# Hypoxia induced carbonic anhydrase mediated dorsal horn sensory neuron activation and induction of neuropathic pain

**DOI:** 10.1101/2021.06.08.447539

**Authors:** M.E. Da Vitoria Lobo, N. Weir, L. Hardowar, Y. Al Ojaimi, R. Madden, Alex Gibson, S.M. Bestall, C Schaffer, M. Hirashima, L.F Donaldson, D.O. Bates, R.P. Hulse

**Affiliations:** Division of Cancer and Stem Cells, School of Medicine, Biodiscovery Institute, University of Nottingham, Nottingham NG7 2UH; School of Science and Technology, Nottingham Trent University, Nottingham, NG11 8NS; Pain Centre Versus Arthritis and School of Life Sciences, The Medical School QMC, University of Nottingham, Nottingham NG7 2UH; Division of Vascular Biology, Kobe University; Nancy E. and Peter C. Meinig School of Biomedical Engineering, Cornell University; Centre of Membrane and Protein and Receptors (COMPARE), University of Birmingham and University of Nottingham, Midlands, UK

**Keywords:** pain, neuron, endothelial, spinal cord, hypoxia, diabetes

## Abstract

Neuropathic pain such as that seen in diabetes mellitus, results in part from central sensitisation in the spinal cord dorsal horn. However, the mechanisms responsible for such sensitisation remain unclear. There is evidence that disturbances in the integrity of the spinal vascular network can be a causative factor in the development of neuropathic pain. Here we show that reduced blood flow and vascularity of the dorsal horn leads to the onset of neuropathic pain. Using rodent models (type 1 diabetes and an inducible endothelial specific vascular endothelial growth factor receptor 2 knockout mouse) that result in degeneration of the endothelium in the dorsal horn we show that spinal cord vasculopathy results in nociceptive behavioural hypersensitivity. This also results in increased hypoxia in dorsal horn sensory neurons, depicted by increased expression of hypoxia markers hypoxia inducible factor 1*α*, glucose transporter 3 and carbonic anhydrase 7. Furthermore, inducing hypoxia via intrathecal delivery of dimethyloxalylglycine leads to the activation of dorsal horn sensory neurons as well as mechanical and thermal hypersensitivity. This shows that hypoxic signalling induced by reduced vascularity results in increased hypersensitivity and pain. Inhibition of carbonic anhydrase activity, through intraperitoneal injection of acetazolamide, inhibited hypoxia induced pain behaviours. This investigation demonstrates that induction of a hypoxic microenvironment in the dorsal horn, as occurs in diabetes, is an integral process by which sensory neurons are activated to initiate neuropathic pain states. This leads to the conjecture that reversing hypoxia by improving spinal cord microvascular blood flow could reverse or prevent neuropathic pain.

## Introduction

Chronic pain is a significant burden faced by patients, with pain not restricted to a single condition but developing due to an array of differing health and disease related afflictions. Approximately 43% of the United Kingdom (1) and 25 % of the global population (2) suffer from chronic pain, which can arise either idiopathically, as a consequence of another disease (e.g. diabetes(3)), or as a consequence of therapy (e.g. chemotherapy(4)). Neuropathic pain typically is described as long-lasting pain, presented as a hypersensitivity to evoked sensory stimuli (e.g. allodynia and hyperalgesia) as well as ongoing pain (5, 6). There is a severe lack of effective analgesic management in the clinic (7). Chronic pain develops as a result of sensitisation of the somatosensory system, with the modulation of intrinsic sensory neural circuits fundamental to the maintenance of nociception (8). Dorsal horn sensory neurons possess the ability to adapt to neuronal stress through key mechanisms that underlie maladaptive plasticity (9). A fine balance between excitatory and inhibitory processing is typically maintained, but during chronic pain excitatory pathways predominate(10), with GABAergic excitation a contributing factor (11, 12). How these sensory neuronal networks alter in times of stress is unknown.

Neural tissues are highly susceptible to depleted oxygen availability, though can reprogramme metabolic pathways to support continued neuronal function (13). Perturbed vascular function accompanies the development of neurodegenerative disease such as Alzheimers(14), and a causal relationship between peripheral limb ischaemia and onset of pain has been shown(15, 16). Previous work has explored systems level modulation of blood flow and the resulting impact upon nociceptive processing and modulation of pain behaviours (17, 18). We have previously shown that a reduction of dorsal horn spinal cord blood flow in a rodent model of diabetes, potentially due to vascular rarefaction, was associated with the onset of pain hypersensitivity (17). Vascular integrity and survival is mediated by the VEGF family of proteins, signalling through VEGFR2. However, a systemic endothelial specific knockout of VEGFR2, which would be predicted to reduce vascular density in the spinal cord, did not induce pain hypersensitivity, and reduced sensitivity in an animal model of arthritis, by reducing spinal cord microglial activation in response to the inflammatory stimulus(18). We therefore set out to determine whether vascular rarefaction in the spinal cord *per se* was able to initiate neuropathic pain phenotypes.

Here we show that the spinal cord microvasculature is critical in maintaining sensory neuronal modulation and demonstrate that alterations in vascular support to the dorsal horn contribute to the development of diabetic neuropathic pain. We use type I diabetic and transgenic rodent models (endothelial specific knockout of vascular endothelial growth factor receptor 2 (VEGFR2) knockout) of reduced spinal cord vascularity to demonstrate that localised vascular degeneration induces a hypoxic microenvironment in the dorsal horn, leading to activation of sensory neurons and initiation of neuropathic pain. The initiation of neuropathic pain is dependent on the response to hypoxia, in particular the upregulation of hypoxia inducible genes and the resultant change in bicarbonate metabolism, which can be modified by inhibitors of the carbonic anhydrase pathway. These results show a route to a potential new therapeutic strategy for analgesics for chronic pain states.

## Methods

### Ethical Approval and Animals used

All experiments using animals were performed in accordance with the UK Home office animals (Scientific procedures) Act 1986 and EU Directive 2010/63/EU. Experimental design and procedures were considered in line with the ARRIVE guidelines, with review by the local Animal Welfare and Ethics Review Boards (University of Nottingham and Nottingham Trent University). These Animals had *ad libitum* access to standard chow and were housed in groups under 12:12h light:dark conditions.

### Transgenic Mice and Chemical Administration

Tie2CreER^T2^ mice (European Mutant Mouse Archive Tg(Tek-cre/ER^T2^)1Arnd,) were interbred with homozygous *vegfr2*^fl/fl^ mice (as previously described (18-20)). This generates tamoxifen inducible endothelial specific vegfr2 knockout adult mice. Mice (both genders) used were vegfr2^fl/fl^ and were either Tie2CreER^T2^ negative (n=83, termed WT) or positive (n=83, termed VEGFR2^CreERT2^). Mice were treated with either intraperitoneal injection of 1mg tamoxifen per mouse (10% ethanol in sunflower oil) once daily for 5 days consecutively (n=116) (17, 18) referred to as VEGFR2^ECKO^ or, a single intrathecal injection of 1µM 4-hydroxytamoxifen (OHT; 10% ethanol in sunflower oil) (n=34) referred to as VEGFR2^scECKO^. All transgenic animals were taken to 8 days post first tamoxifen or OHT injection. All intrathecal injections were performed under brief isoflurane anaesthesia (∼2% oxygen).

For the hypoxia mimetic study, 1mM Dimethyloxalylglycine (DMOG, in PBS, Tocris) or vehicle was intrathecally injected into C57Bl6 adult male (35 mice) under recovery anaesthesia (isoflurane ∼2% oxygen). Acetazolamide was delivered via intraperitoneal injection (40 mg/kg, in 1%DMSO in PBS, Tocris). Db/db (10) mice were compared WT lean (10) controls..

### Streptozotocin induced Diabetes

Sprague Dawley female rats (∼250g) were used and treated via intraperitoneal injection with either vehicle (citrate buffer; n=6) or Streptozotocin (in citrate buffer, 50mg/kg; STZ). (Hulse et at. 2015). Hyperglycaemia was determined as >15mmol in samples of blood extracted via tail prick at 1 week post STZ injection. Under isoflurane anaesthesia (∼2% oxygen), one third of an insulin pellet (LinShin,Canada) was implanted in the scruff of the neck using a trocar, in STZ treated animals. Experimental group animal body weights and blood glucose measurements were recorded during the study. Measures (mean ±SEM) recorded at the end of the study (week 8) are presented below; Blood glucose (mmol/dL): naïve=7.44±0.29, diabetic =28.5±1.82, Animal body weight(gms): naïve=330.6±4.49, diabetic=284.2±4.86.

### Nociceptive Behaviour

Animals were habituated to the testing environment prior to nociceptive behavioural experimentation, which were performed as previously described (17). Mechanical withdrawal thresholds were determined via application of von Frey (vF) monofilaments to the hindpaw plantar surface. vF filaments of increasing diameter were applied (a maximum of five seconds or until paw withdrawal), with each vF filament applied a total of 5 times to generate force response curves. Withdrawal frequencies of 50% at each weight was used to determine the mechanical withdrawal threshold (g)(21). The Hargreaves test was performed to measure thermal nociceptive behaviour(22). A radiant heat source was focused on the plantar surface of the hindpaw. The time taken for the mouse to withdraw their paws were recorded. The latency to hindpaw response was measured 3 times. For motor behavioural assessment, mice were placed in Perspex enclosures and were left undisturbed for 5 minutes whilst being video recorded to explore the environment (23). Following experimentation animals were returned to housing cages and data analysed offline.

### Immunofluorescence

Animals were terminally anaesthetised with sodium pentobarbital (60mg/kg, Sigma-Aldrich), and transcardially perfused with PBS followed by 4% PFA (pH7.4). Tissue was dissected, cryoprotected (30% sucrose) and stored at −80°C until spinal cords were cryosectioned (40µm thickness) (Ved et al., 2018). Slides were incubated in primary antibodies (see table 1) in blocking solution (5% bovine serum albumin, 10% fetal calf serum, 0.2% Triton-X) at 4°C for 72 hours, then washed in PBS. Slides were subsequently incubated in secondary antibodies in PBS + 0.2% Triton X-100 at room temperature for 2 hours. Confocal microscopy of the dorsal horn of the lumbar region of the spinal cord was performed using a Leica SPE confocal microscope.

**Table 1.**
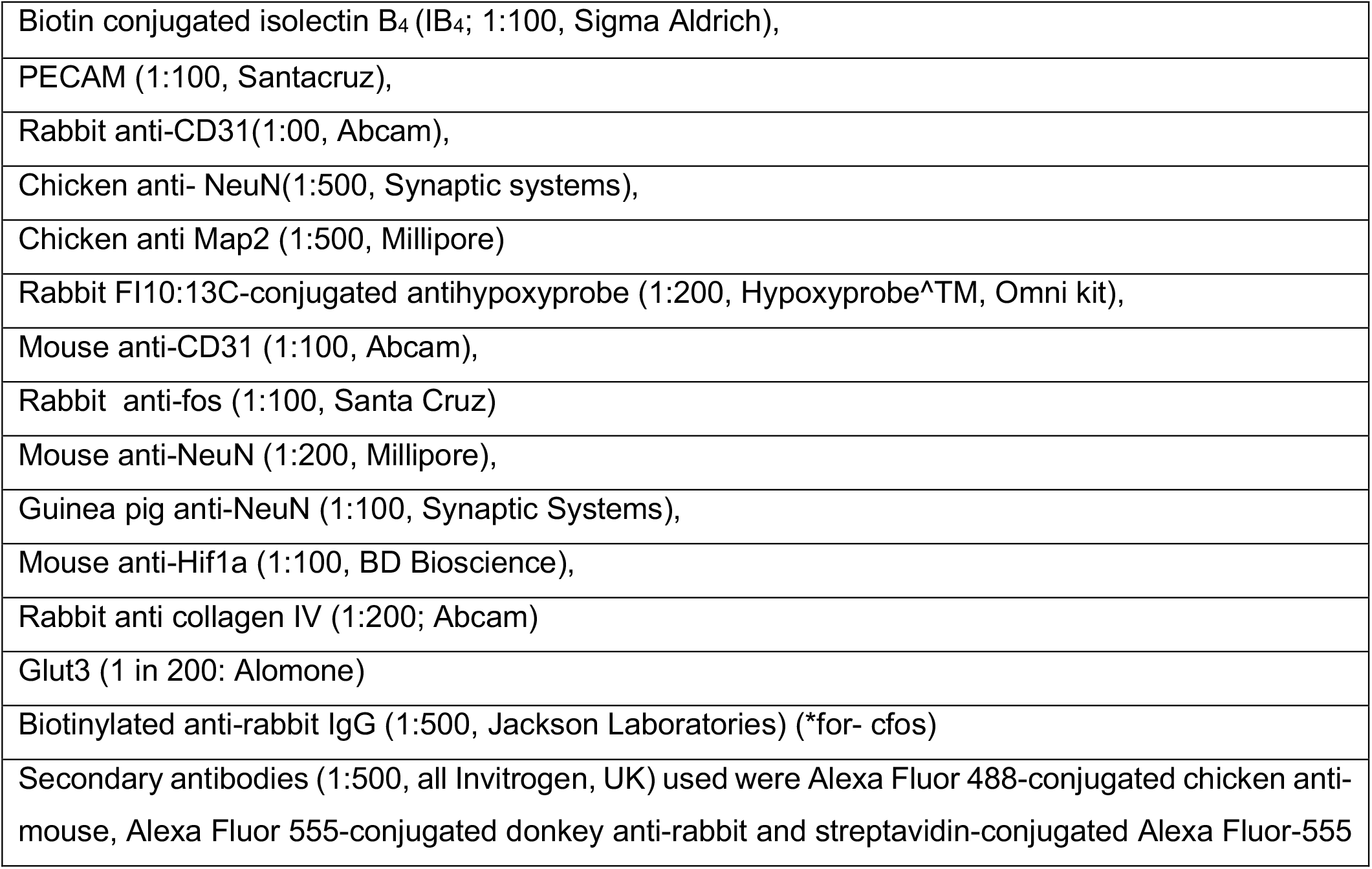
– List of Primary and Secondary Antibodies for Immunofluorescence

**Table 2.**
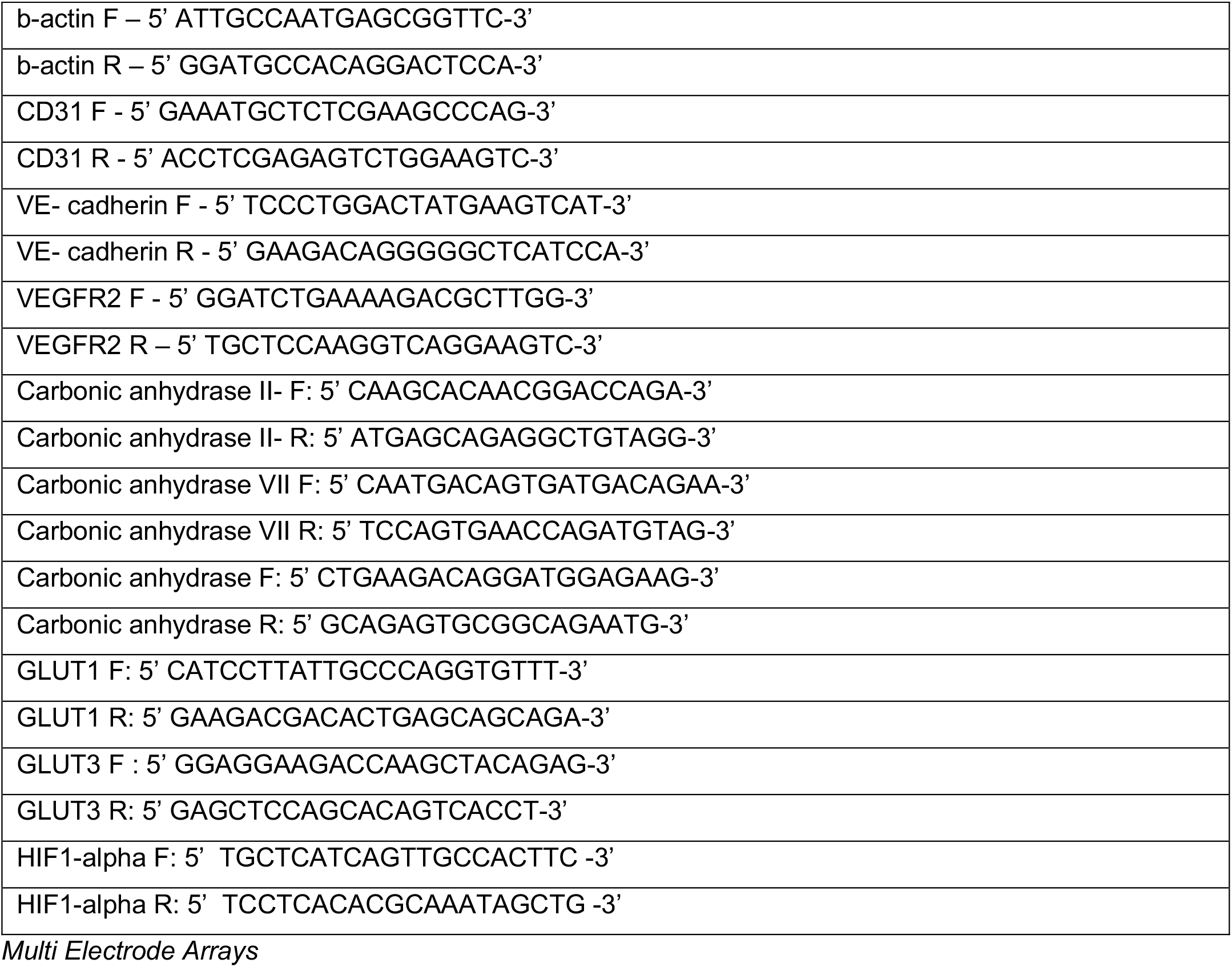
– List of Primers Used

### Hypoxyprobe Assay

Hypoxyprobe staining was performed as outlined in the manufacturers instructions (Hypoxyprobe omni kit). Transgenic mice were used 8 days post intrathecal injection of hydroxytamoxifen. Experimental animals were administered 60mg/kg hypoxyprobe via intraperitoneal injection. Animals were terminally anaesthetised with sodium pentobarbital (60mg/kg, Sigma-Aldrich) 30 minutes later and transcardially perfused with PBS followed by 4% paraformaldehyde (PFA; pH7.4). Lumbar spinal cords were extracted and cryosectioned as outlined above. Anti-pimonidazole rabbit antisera; (PAb2627AP) was added to slides at 1 in 200 dilution overnight at 4°C and subsequently incubated in Alexafluor-555-conjugated donkey anti-rabbit in PBS + 0.2% Triton X-100 at room temperature for 2hrs. Confocal imaging of the dorsal horn of the lumbar spinal cord was performed on a Leica TCS SPE confocal microscope.

### TRITC dextran perfusion Assay

Transgenic mice (8 days post administration of hydroxytamoxifen) were terminally anaesthetised and anaesthesia was maintained with sodium pentobarbital (60mg/kg, Sigma-Aldrich). The external jugular vein was cannulated and 1 mg per 20g tetramethylrhodamine isothiocyanate (TRITC) dextran (76 *M*^w^; Sigma-Aldrich) was injected (24, 25). Mice were monitored for 30 minutes post TRITC dextran infusion. Subsequently animals were transcardially perfused with 4% PFA (pH7.4) and lumbar spinal cords were extracted and samples prepared for immunofluorescent confocal imaging as outlined previously.

### Intravital imaging

C57Bl6 male mice (∼30g) were deeply terminally anaesthetised and anaesthesia maintained (ip Ketamine, medetomidine). Body temperature was maintained (∼ 37°C) using a feedback-controlled heating blanket with a rectal probe. Midline incision was made along the thoracic-lumbar region of the mouse and muscle was dissected from the vertebrae. A laminectomy was performed between thoracic (T10) and lumbar vertebra (L5) to expose the lumbar spinal cord. In house stainless steel spinal vertebrae clamps were used to stabilised the vertebrae whilst an in house window chamber was attached to the spinal cord vertebra (26) The spinal cord hydration was maintained with physiological saline prior to the application of Kwik-Sil® silicone elastomer (World Precision Instruments) and a 5mm coverslip number 1 (Harvard Apparatus). Following this the mouse was placed on to the confocal microscope (Leica SPE) and images were captured via 10x objective. Mice were administered alexafluor 555 conjugated Wheat germ agglutinin (1mg/ml, Thermoscientific) via tail vein injection. Upon identification of the endothelium, sodium fluorescein (i.p. 100mg/ml) was administered intraperitoneally (27). Image acquisition began immediately following sodium fluorescein injection. Spinal cords were treated with either vehicle (PBS) or 2.5nM VEGF-A^165^a with or without VEGFR2 inhibitor ZM323881 (100nM) for 5 minutes prior to fluorescein injection (as described by (28)). Sodium fluorescein permeability was measured as previously described in the mesentery (29) that involved determining a region of interest extending across the vessel width and encompassing the region outside the vessel wall with solute permeability determined.

### Spinal Cord Endothelial cell culture

Spinal cord endothelial cells were isolated as described in (Ved et al., 2018). Whole lumbar spinal cords were dissected from C57Bl6 mice and incubated with 1.25% collagenase in endothelial cell media (M199 media, 60mg/ml endothelial cell growth supplement and 50ug/ml Heparin; Sigma-Aldrich) for 30 minutes at 37°C, 5% oxygen. The cell suspension was centrifuged at 240g for 5 minutes. The supernatant was discarded and the pellet was resuspended in media with an equal volume of 1% BSA and centrifuged at 1200g for 20 minutes. The BSA/myelin fraction was aspirated and discarded. The remaining cell pellet resuspended in endothelial cell media and plated onto a 96-well sterile plates coated with 1% gelatin. Upon 80% cell confluency, spinal cord endothelial cells were treated for 24hrs with either media supplemented with either vehicle (10% ethanol in sunflower oil) or OHT. Cell viability was determined with Neutral red (Sigma-Aldrich).

### Spinal Isolated Neuronal Cell Culture

Spinal cord neurons were dissected as described by (Brewer and Torricelli, 2007). Whole spinal cords were isolated from C57Bl6 mice and digested in papain (2mg/ml, Worthington, cat. no. 3119) for 30 minutes at 30°C. The cell suspension was triturated using Pasteur pipette. To purify neurons a density gradient separation preparation was performed using OptiPrep (Sigma Aldrich) density gradient. The neuronal containing fraction was collected and centrifuged (1100 rpm 2 minutes) to obtain a pellet. The pellet was resuspended in Neurobasal media (supplemented with B27 and L-Glutamine, Invitrogen) and plated on sterile 6 or 24-well plates coated with poly-D-Lysine (sterile lyophilized 135 kD; Sigma) and 1mg/ml laminin. Cells were exposed to hypoxia (1% O^2^) or normoxia for 24 hours.

### Immunocytochemistry

Spinal cord neurons were isolated as outlined and plated on glass coverslips coated in poly-l-lysine (24hrs, Sigma Aldrich) followed by laminin (5mg/ml 2hrs, Sigma Aldrich). They were cultured in media for 24hrs and subsequently washed in PBS, prior to incubation in 4% PFA for 15 minutes at room temperature. Neurons were permeabilised with 0.4% Triton X-100 in PBS for 15 minutes at room temperature. Cells were incubated in blocking solution (5% bovine serum albumin, 10% fetal calf serum, 0.2% Triton-X), mixed gently and incubated for 30 min at room temperature. Cells were incubated in primary antibodies for identification of neurons (chicken anti-MAP2, SySy) overnight at 4°C. Cells were subsequently washed in PBS and secondary antibodies (Alexa Fluor 555-conjugated donkey anti-mouse, Alexa Fluor 488-conjugated donkey anti-chicken) added for 2hrs at room temperature. Glass coverslips were mounted on slides using Vector Shield (H-1000, Vector Labs).

### qPCR Method

Whole lumbar spinal cord lysates and isolated lumbar spinal cord neurons were used to extract RNA in experimental groups using TRIzol reagent (Invitrogen). 1ug RNA was reverse transcribed to cDNA using PrimeScript™RT reagent kit (TaKaRa, RR037A). Quantitative PCR was done using a LightCycler 480 SYBR Green I Mastermix (Roche 04707516001) following the manufacturer’s instructions. CD31 and VE-cadherin primers were obtained from Eurofins. All other primers were obtained from Sigma-Aldrich. The reference gene used was b-actin. The primer sequences were as follows (5’-3’);

### Multi Electrode Arrays

Mouse lumbar spinal cord were extracted from five C57.Bl6 mice and maintained in neurobasal media (supplemented with B27 and L-Glutamine, Invitrogen). Lumbar spinal cord tissue was cut to 400μm thickness and incubated on high density microelectrode arrays (3Brain, 4096 electrodes). Neuronal electrical activity was recorded digitally via the Biocam X after 24hrs treatment with either vehicle or 1mM DMOG application. Data was recorded offline using Brainwave analysis software.

### Western Blotting

Protein was extracted from lumbar spinal cord tissue and L3-L5 dorsal root ganglion as previously described (17, 21) from the animals from the experimental groups (VEGFR2KO and intrathecal DMOG studies compared to vehicle experimental controls) as above. In additional experiments, protein lysate samples were also extracted from invitro lumbar spinal cord slices prepared from six C57bl6 male adult mice and slices maintained in Neurobasal media (supplemented with L-Glutamine, Invitrogen)) and exposed to vehicle or 1mM DMOG for 24 hours. Tissue was lysed using RIPA buffer (ThermoFisher) containing protease inhibitor cocktail (20 µl ml−1 buffer; 1 mM phenylmethylsulfonyl fluoride, 10 mM sodium orthovanadate and 50 mM sodium fluoride, Sigma Aldrich). Equal protein lysate concentrations were loaded on a 4%-20% precast Mini-Protean gradient TGX gel (Biorad), separated by SDS-PAGE electrophoresis and transferred using Trans-blot turbo transfer system (Bio-Rad) to a PVDF membrane. The membrane was incubated in 5% BSA in tris-buffered saline (TBS)-Tween 0.1% (TBST) for 1 hour at room temperature, followed by incubation with primary antibodies (table 3) overnight at 4 °C. The membrane was then incubated in secondary antibodies in TBST-0.1%-1% BSA. Membranes were washes three times with TBST and visualised on the Licor Odyssey Fc.

**Table 1 –.** List of Primary and Secondary Antibodies

|  |
| --- |
| HIF1-alpha anti mouse (1 in 100, BD Bioscience) |
| Actin - Rabbit polyclonal anti- Beta Actin (1 in 100 Santa Cruz) |
| Actin - Goat polyclonal anti- Beta Actin (1 in 200 Santa Cruz) |
| Tubulin – anti-rabbit anti-α tubulin (1 in 1000, Cell Signalling, #2144) |
| ZO-1 - rabbit anti-ZO1 (1 in 500, Invitrogen, Cat #61-7300) |
| Occludin - rabbit anti-occludin (5 μg ml<sup>-1</sup>, Invitrogen) |
| CD-31 mouse monoclonal anti-CD31 (2μg/ml, Abcam; AB24590), |
| CA-7 rabbit (1 in 500, Cell Signalling, # PA5-103703) |
| Vascular Endothelial Growth Factor 2 (VEGFR2) – rabbit anti VEGFR2 (1 in 500, Cell Signalling, #2479) |
| Licor donkey anti-goat, anti-rabbit and anti-mouse antibodies 1:10000 |

### Statistical Analysis

All data are represented as mean±SEM unless stated. Data was acquired/quantified using Microsoft Excel and Graphpad Prism 8 Quantification of immunofluorescence was quantified by obtaining 5 random images from 5 random non-sequential sections (Z stacks) per animal and a mean value calculated per animal. Images were analysed using ImageJ (https://imagej.nih.gov/ij/) or Imaris 8.11 (Bitplane) to determine vessel diameter and volume. Dorsal horn neuronal number was determined according to lamina I-V as previously characterised (30) (Supplemental Figure 1C). Densitometry analysis of western blot was performed using gel quantification plugin in ImageJ. Mean fluorescence intensity was measured over time enabling visualisation of vessel perfusion and permeability. Nociceptive behaviour and the neuron number in the dorsal horn of the spinal cord were analysed using Two way ANOVA with post hoc analysis. Western blot densitometry, QPCR, MEA recordings and tritc dextran perfusion were analysed using a Mann-Whitney test. Video analysis of rodent motor behaviour was performed using MATLAB (23) and analysed using Mann-Whitney test

**Figure 1 –.**
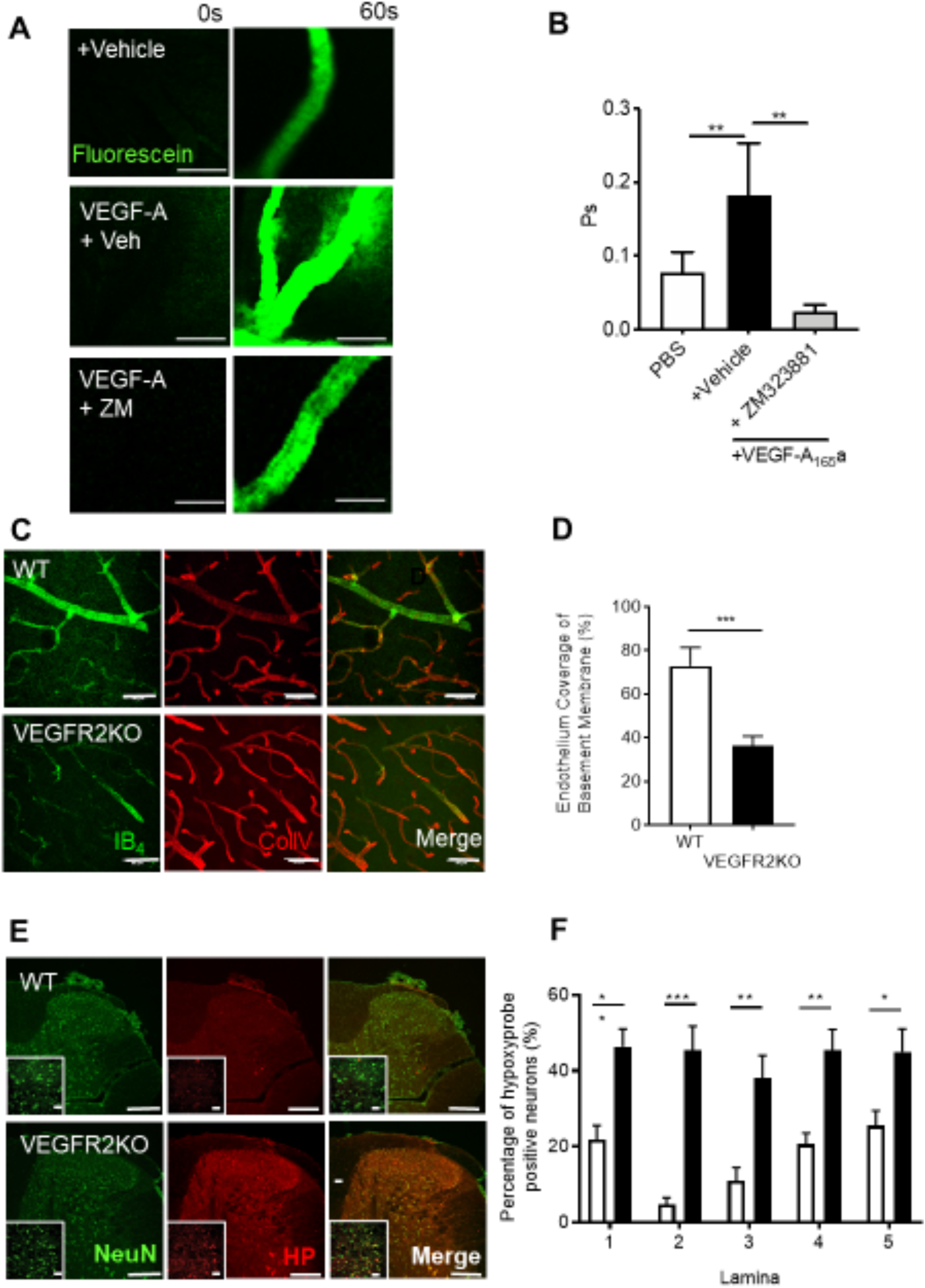
Dorsal horn hypoxia develops in a rodent model of type 1 diabetic neuropathic pain Type 1 diabetes was induced in rats by intraperitoneal injection of streptozotocin supplemented with insulin. Diabetic rsodents developed [A] decreases in mechanical withdrawal threshold and [B] reduced latencies to heat withdrawal compared to age-matched control animals (Naïve). [C] Cryosections of the dorsal horn from Naïve and Diabetic experimental groups were stained for basement membrane (Collagen IV) and endothelium (IB^4^). [D] In the diabetic dorsal horn there was an increased percentage of collagen sleeves (collagen IV positive, endothelium (IB4) negative, arrows) compared to age matched naïve animals (n=5 per group, ** P<0.01 Unpaired T Test). [E] Representative confocal microscopy images of hypoxyprobe (HP) positive sensory neurons (NeuN Labelled) in the dorsal horn of the spinal cord in naïve and diabetic cryosections (co-labelled with NeuN, low power images (main) scale bar = 100μm, high power images (inset) = 50 μm). [F] Mapping of the dorsal horn laminae demonstrates increased hypoxyprobe positive neurons/lamina in diabetic animals when compared with naïve control animals (*P<0.05, ***P<0.001 Two Way ANOVA with Tukey’s multiple comparison, n=5 per group).

## Results

### DorsalHornHypoxiaInduction

To determine whether increased diabetic neuropathic pain was associated with vascular loss and hypoxia, we used a rat model of type 1 (streptozotocin induced and insulin supplemented) that led to the development of mechanical (Fig. 1A) and heat (Fig. 1B) nociceptive behavioural hypersensitivity. This was associated with an increase in collagen positive microvessels, negative for IB^4^ labelled endothelium (collagen sleeves) in diabetic animals compared to age matched vehicle control animals (Fig. 1C and 1D) indicative of loss of endothelium and vascular rarefaction. Additionally, there was a significant increase in the number of hypoxic dorsal horn neurons (labelled with hypoxyprobe) in diabetic animals when compared with age matched vehicle control animals (Fig. 1E and 1F, Supplemental Figure 1A), and a small decrease in the total number of identified neurons in the dorsal horn (NeuN labelled; Supplemental Figure 1B and C). These results indicate that hypoxia caused by loss of blood vessels could play a part in induction of diabetic neuropathic pain.

We have previously shown that VEGFR2^ECKO^ animals had no change in nociceptive processing compared with wild type animals and that they were actually resistant to hyperalgesia and allodynia induced by induction of arthritis. To determine whether these mice had a change in vessel density, hypoxia and cell number, we stained the spinal cord of these animals after induction with tamoxifen and knockout. VEGFR2^ECKO^ animals showed an increase in the number of collagen sleeves compared with age matched litter mate control wild type animals (Fig. 2A and 2B), indicating vascular rarefaction. Although VEGFR2^ECKO^ did not affect the number of neurons (NeuN labelled) in the dorsal horn across lamina 1-5 (Supplemental Figure 2A-D). had significantly increased the number of hypoxyprobe labelled neurons in the dorsal horn compared with control animals (Fig. 2C and 2D, Supplemental Figure 2E) demonstrating hypoxia as a consequence of VEGFR2^ECKO^.

**Figure 2 –.**
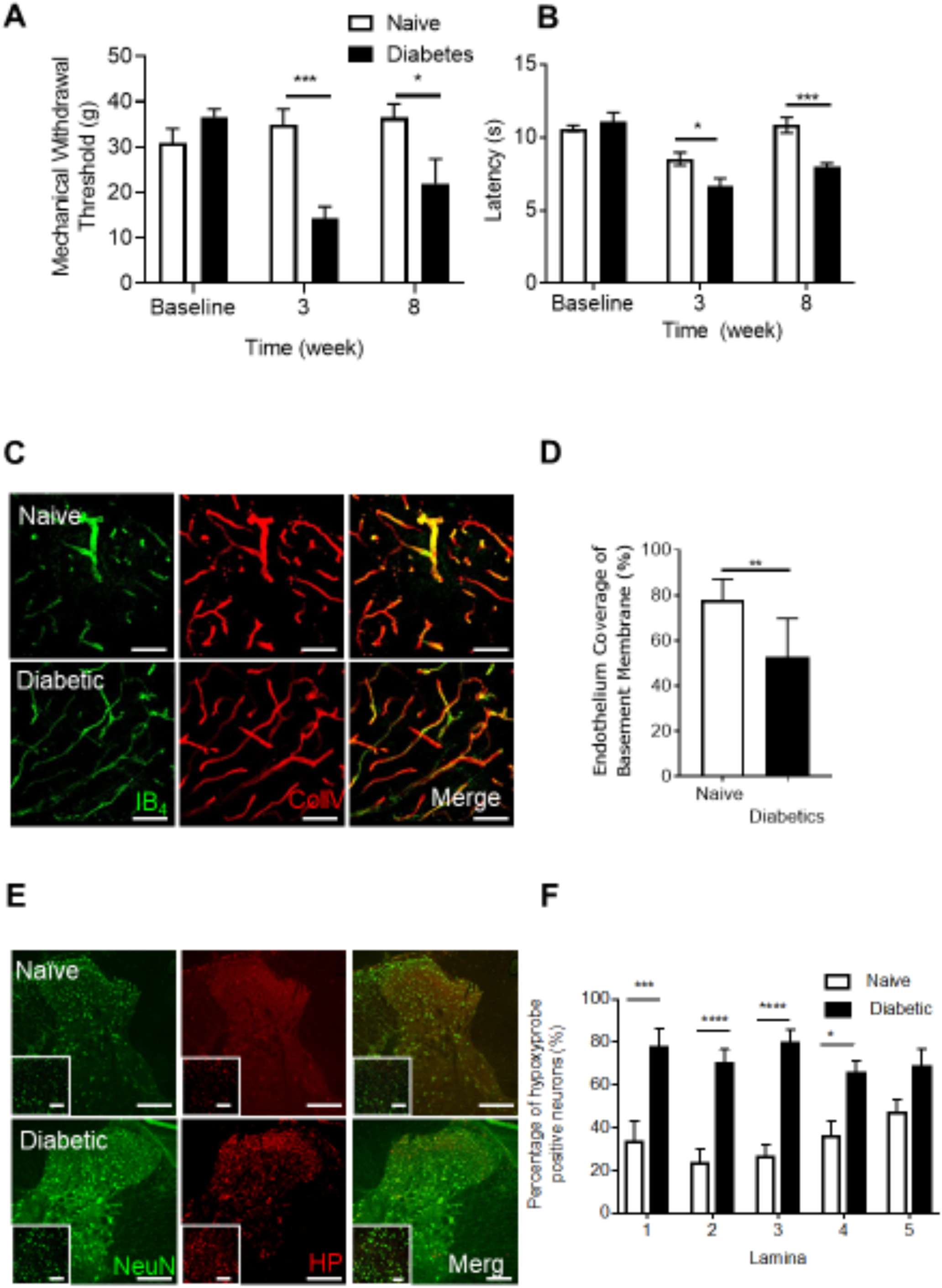
Vascular endothelial growth factor modulation of the spinal cord blood vasculature [A] Dorsal horn cryosections were imaged via confocal microscopy to identify microvessels (Collagen IV -basement membrane and IB^4^ – endothelium, scale bar = 50 μm) in WT and VEGFR2^ECKO^ mice. [D] In VEGFR2^ECKO^ mice there was an increased percentage of collagen sleeves (microvessels collagen IV positive endothelium (IB4) negative) compared to WT mice (n=5 per group, ** P<0.01 Unpaired T Test). Both WT and VEGFR2^ECKO^ mice were treated via intraperitoneal injection of hypoxyprobe. [E] Confocal microscopy of the dorsal horn was used to identify hypoxyprobe (HP) labelled sensory neurons (co-labelled with NeuN, low power images (main) scale bar = 100μm, high power images (inset) = 50 μm)) in WT and VEGFR2^ECKO^ mice. [F] VEGFR2^ECKO^ mice had increased percentage of hypoxic sensory neurons across dorsal horn laminae 1-5 compared to WT mice (*P<0.05, **P<0.01, ***P<0.001 Two Way ANOVA with Tukey’s multiple comparison, n=5 per group). [G] Top confirm VEGFR2 signalling in the spinal cord, microvessels were identified in the dorsal horn of the lumbar region of the spinal cord in C57 Bl6 mice. 100mg/ml sodium fluorescein was administered via intraperitoneal injection, with data acquired via time lapse confocal imaging allowing for recording of solute permeability (Ps, scale bar = 20 μm). Representative images of fluorescein filled microvessels in the spinal cord from experimental groups -PBS control and Vascular Endothelial Growth Factor-A^165^a (VEGF-A^65^a, 2.5nM) with either vehicle (0.01% DMSO in saline) or VEGFR2 specific antagonist ZM323221 (100 nM). [H] VEGF-A^65^a treatment led to increased fluorescein leakage from the dorsal horn microvessels (**P<0.01 One Way ANOVA with Tukey’s multiple comparison, PBS n = 5, VEGF-A^65^a +vehicle n = 6). Treatment with ZM323881 attenuated VEGF-A^165^ induced vascular leakage (**P<0.01 One Way ANOVA with Tukey’s multiple comparison, VEGF-A^165^ + ZM323881 n = 7).

Blood vessel survival has previously been shown to require endogenous VEGFR2 activation in tissues including the oesophagous, skin and salivary gland (Kamba et al 2006) as well as in other tissues (Lee et al). However, it is not know whether VEGFR2 signalling is required for vessels in the spinal cord, which form part of the blood brain barrier. We therefore sought to determine whether spinal cord vascular endothelial growth factor 2 (VEGFR2) signalling is intact in spinal cord microvessels. VEGF-A^165^a was administered directly to the dorsal horn of the spinal cord, and vascular permeability measured using intravital imaging. VEGF-A^165^ increased solute permeability of the blood spinal cord barrier when compared to vehicle control (Fig.2E. and 2F) which was blocked by treatment with VEGFR2 inhibitor ZM323881 (Fig.2E. and 2F).

### Onset of Chronic Pain due to the Induction of Spinal Cord restricted Dorsal Horn Vasculopathy

The previous explored experimental models were systemic rodent models of vasculopathy. Here we developed a refined model that induced vasculopathy restricted to the spinal cord. Endothelial cells (CD31 positive) were extracted from WT and VEGFR2^CreERT2^ mice spinal cords (Fig. 3A). Cell viability was determined following OHT treatment. In a concentration dependent manner OHT treatment led to a significant reduction in endothelial cell viability in VEGFR2^CreERT2^ mice spinal cord endothelial cells when compared with WT mice spinal cord endothelial cells (Fig. 3B). To determine whether this was also true in vivo, spinal cord endothelial specific VEGFR2 knockout (VEGFR2^scECKO^) mice were generated by intrathecal injection of OHT into the lumbar enlargement. This was confirmed by a reduction in VEGFR2 expression (Fig. 3C. immunofluorescence and Fig. 3D QPCR quantification) in the spinal cord of VEGFR2^scECKO^ mice when compared to WT mice. In addition, endothelial markers VE-Cadherin (Fig. 3E.), CD31 (Fig. 3F) and Glucose transporter 1 (GLUT1, Fig. 3F) were also reduced in the spinal cord of VEGFR2^scECKO^ mice when compared with WT mice. Furthermore, VEGFR2^scECKO^ mice had reduced protein expression of endothelial markers (Representative Western Blot, Fig. 3H and quantification of ZO1(Fig. 3I), CD31(Fig. 3J) and occluding (Fig. 3K)) in the spinal cord. Notably, there was no change in protein expression of these endothelial markers (ZO-1 and CD31) in the lumbar 3-5 dorsal root ganglia of either WT or VEGFR2^scECKO^ mice, showing that this induction was limited to the spinal cord (Supplemental Figure 3).

**Figure 3 –.**
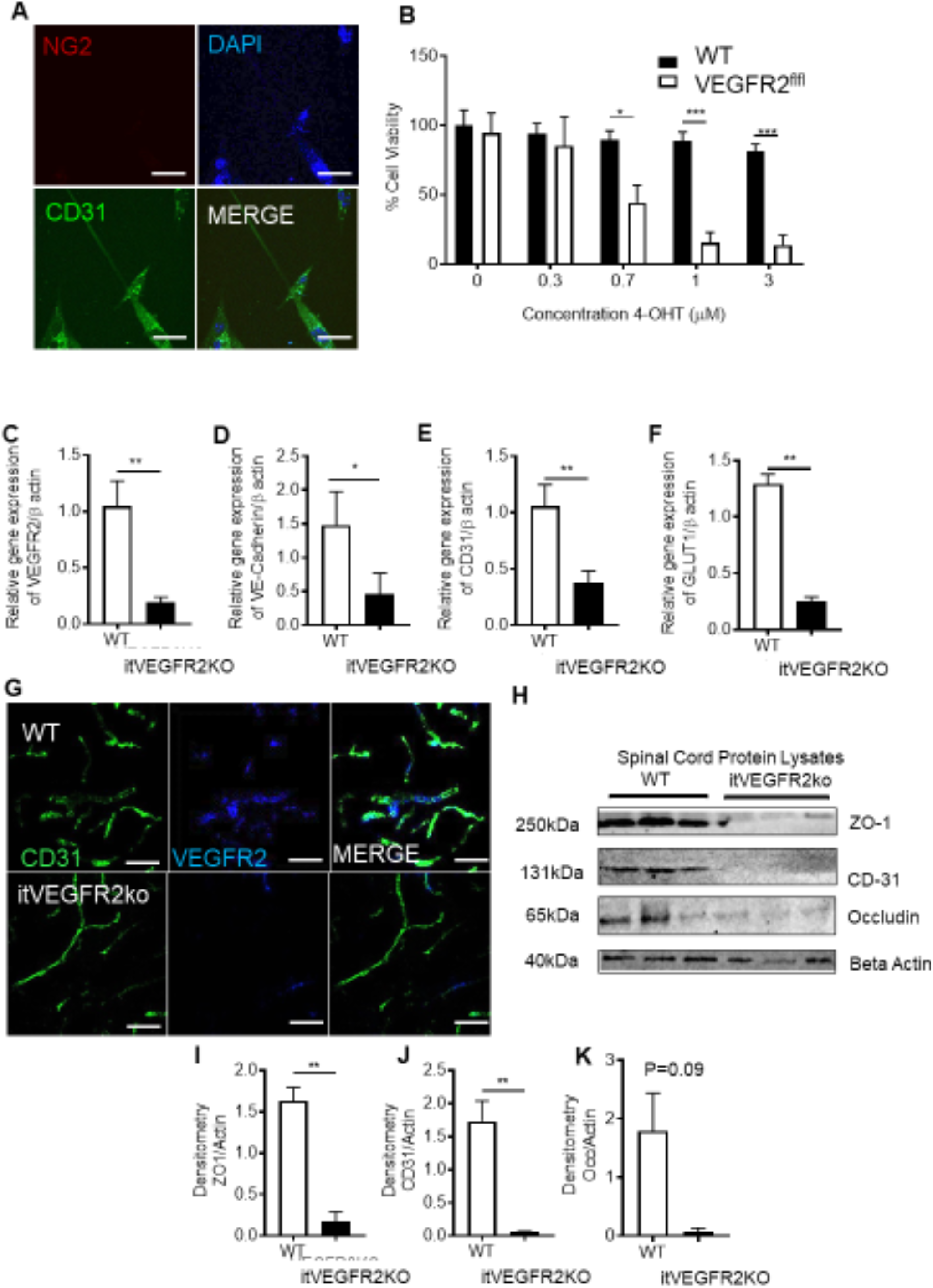
VEGFR2 knockout in the spinal cord induces localized vasculopathy [A] Spinal cord endothelial cells (NG2^-ve^, CD31^+ve^) were isolated from WT and VEGFR2^CreERT2^ transgenic mice (scale bar = 10μm) and [B] cell viability was determined following treatment with OHT. Endothelial cells extracted from WT mice were unaffected by OHT treatment whereas those from VEGFR2^CreERT2^ mice demonstrated a concentration dependent reduction in cell viability(*P<0.05, ***P<0.001 Two Way ANOVA with Tukey’s multiple comparison, n=3 per group). [C] Intrathecal injection of 1μM OHT into the lumbar region of the spinal cord of VEGFR2^CreERT2^ mice (VEGFR2^scECKO^) led to reduced VEGFR2 mRNA expression in the lumbar spinal cord compared with similarly treated WT mice. There were reductions in mRNA expression of endothelial specific markers [D] VE-Cadherin, [E] CD31 and [F] GLUT1 (*P<0.05, **P<0.01, Unpaired T Test, n=5 per group). [G] VEGFR2 immunoreactivity was reduced in CD31 labelled microvessels (scale bar = 50μm) in the dorsal horn of VEGFR2^scECKO^ mice. [H] Representative immunoblots show reductions in protein expression of endothelial markers in the lumbar region of the spinal cord from VEGFR2^scECKO^ mice. Densitometric quantification demonstrates reduced protein expression of [I] ZO1, [J] CD31 and [K] Occludin in VEGFR2^scECKO^ mice (**P<0.01, Unpaired T Test, n=5 per group).

Intrathecal OHT treatment led to significant vascular degeneration (Fig. 4A) of the microvasculature in the dorsal horn of VEGFR2^scECKO^ mice, shown as reduced vessel diameter (Fig. 4B) and decreased volume of total vessel coverage (Fig. 4C). In addition, there was a reduction in dorsal horn vessel perfusion in VEGFR2^scECKO^ mice indicated by decreased TRITC-Dextran labelled vessels (Fig.4D). This reduction in vessel perfusion was accompanied by an increase in hypoxyprobe labelled neurons (Fig. 4F, hypoxyprobe labelled with NeuN) in laminae 1-5 in the dorsal horn (Fig. 4G). There were no alterations in total neuron number in the dorsal horn in either WT or VEGFR2^scECKO^ mice (Supplemental Figure 4A). VEGFR2^scECKO^ mice had heat hypersensitivity (Fig. 4H), but no alterations in mechanical withdrawal thresholds (Fig. 4I) or motor behaviour (Supplemental Figure 4C). No change in heat withdrawal latency or mechanical withdrawal threshold was identified in WT mice following OHT treatment (Fig. 4H and 4I).

**Figure 4 –.**
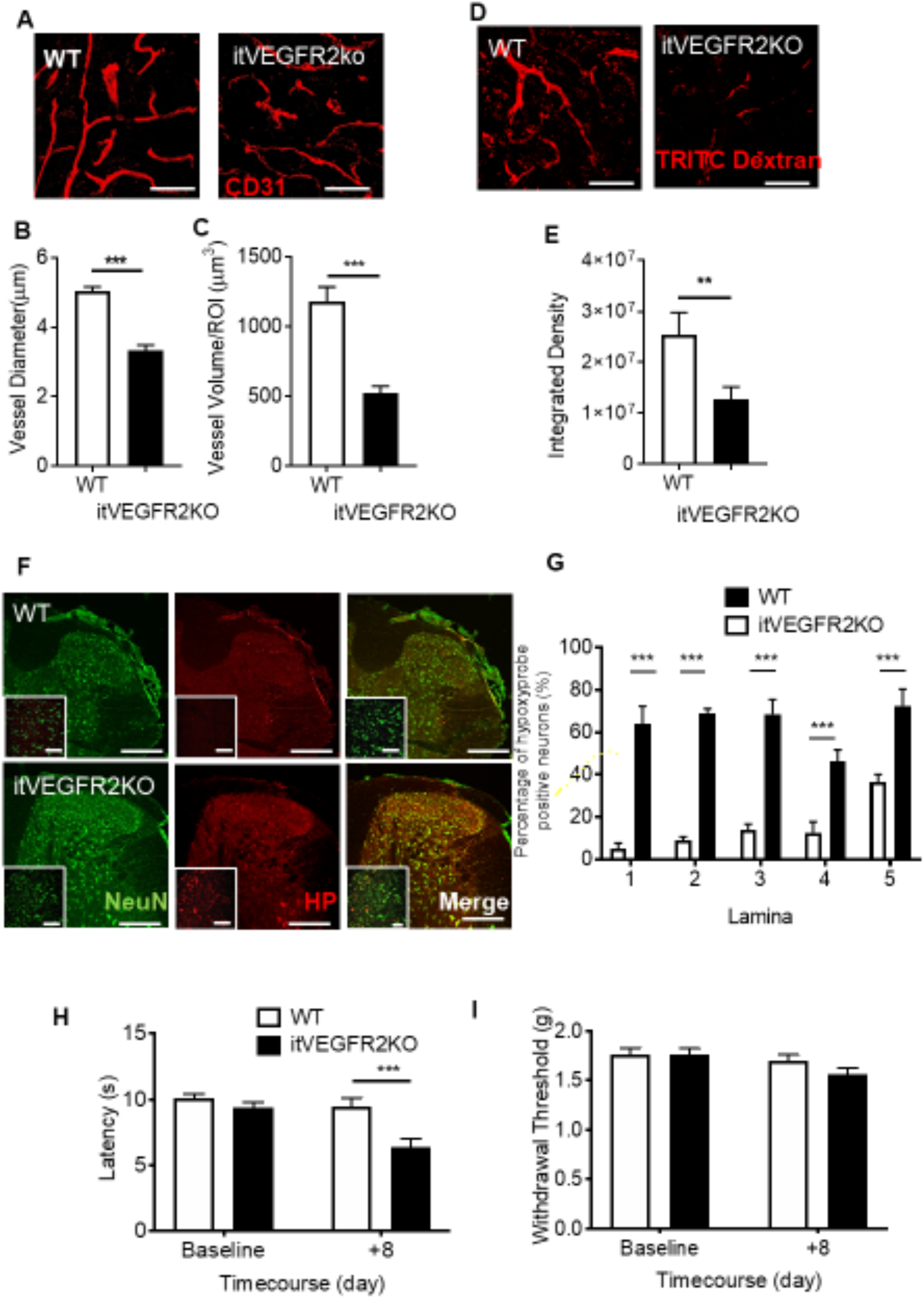
Spinal cord VEGFR2 knockout induced vasculopathy causes hypoxia and chronic pain [A] VEGFR2^scECKO^ mice have degeneration of the dorsal horn endothelium 8 days following injection highlighted by the representative confocal images of the CD31 labelled endothelium (scale bar = 50 μm). OHT treatment led to [B] reduced vessel diameter and [C] decreased vessel volume in VEGFR2^scECKO^ mice (*P<0.05, ***P<0.001, Unpaired T Test, n= 4 per group). [D] TRITC dextran was intravenously injected into terminally anaesthetized WT and VEGFR2^scECKO^ mice and TRITC Dextran positive perfused vessels identified as outlined in the representative confocal images (scale bar = 50 μm). [E] In VEGFR2^scECKO^ mice there was less TRITC Dextran in the dorsal horn compared with WT mice (**P<0.01, Unpaired T Test, n= 5 per group). [F] Representative confocal images of Hypoxyprobe labelled neurons (co-labelled with NeuN) in the dorsal horn of WT and VEGFR2^scECKO^ mice (low power images (main) scale bar = 100μm, high power images (inset) = 50 μm). [G] There was an increased percentage of neurons positive for hypoxyprobe, in each laminae 1-5 across the dorsal horn in VEGFR2^scECKO^ mice (***P<0.001 Two Way ANOVA with Tukey’s multiple comparison, n=5 per group). [H] VEGFR2^scECKO^ mice developed reduced withdrawal latencies to heat (***P<0.001 Two Way ANOVA with Tukey’s multiple comparison, n=16 per experimental group). [I] No change to mechanical withdrawal threshold was observed (n=16 per experimental group).

### Induction of Hypoxia Signalling in the Dorsal Horn Sensory Neurons

VEGFR2^scECKO^ mice demonstrated an increase in expression of Hypoxia Inducible Factor 1 *α* (HIF1*α*) and Glucose Transporter 3 (GLUT3, Supplemental Figure 5A and B) in dorsal horn sensory neurons (colabelled with Neuronal marker NeuN, Fig. 5 A and 5B; Supplemental Figure 5C and D) and a significant increase in HIF1*α* mRNA (Fig. 5C) and that of hypoxia responsive genes Carbonic Anhydrase VII (CA7, Fig. 5D) and GLUT3(Fig. 5E). In isolated neurons from the lumbar region of the spinal cord (Supplemental Figure 6A) exposed to differing environmental oxygen conditions; normoxia and hypoxia (1%) resulted in an increase HIF1*α*, CA7 and GLUT3 (Supplemental Figure 6B). In addition, KCC2 was downregulated (Supplemental Figure 6B). Interestingly, increased HIF1*α* was also increased in a rodent type 1 (STZ induced, Fig 5F-I) and type 2 (db/db obese mice, Fig 5J) models of diabetes.

**Figure 5 –.**
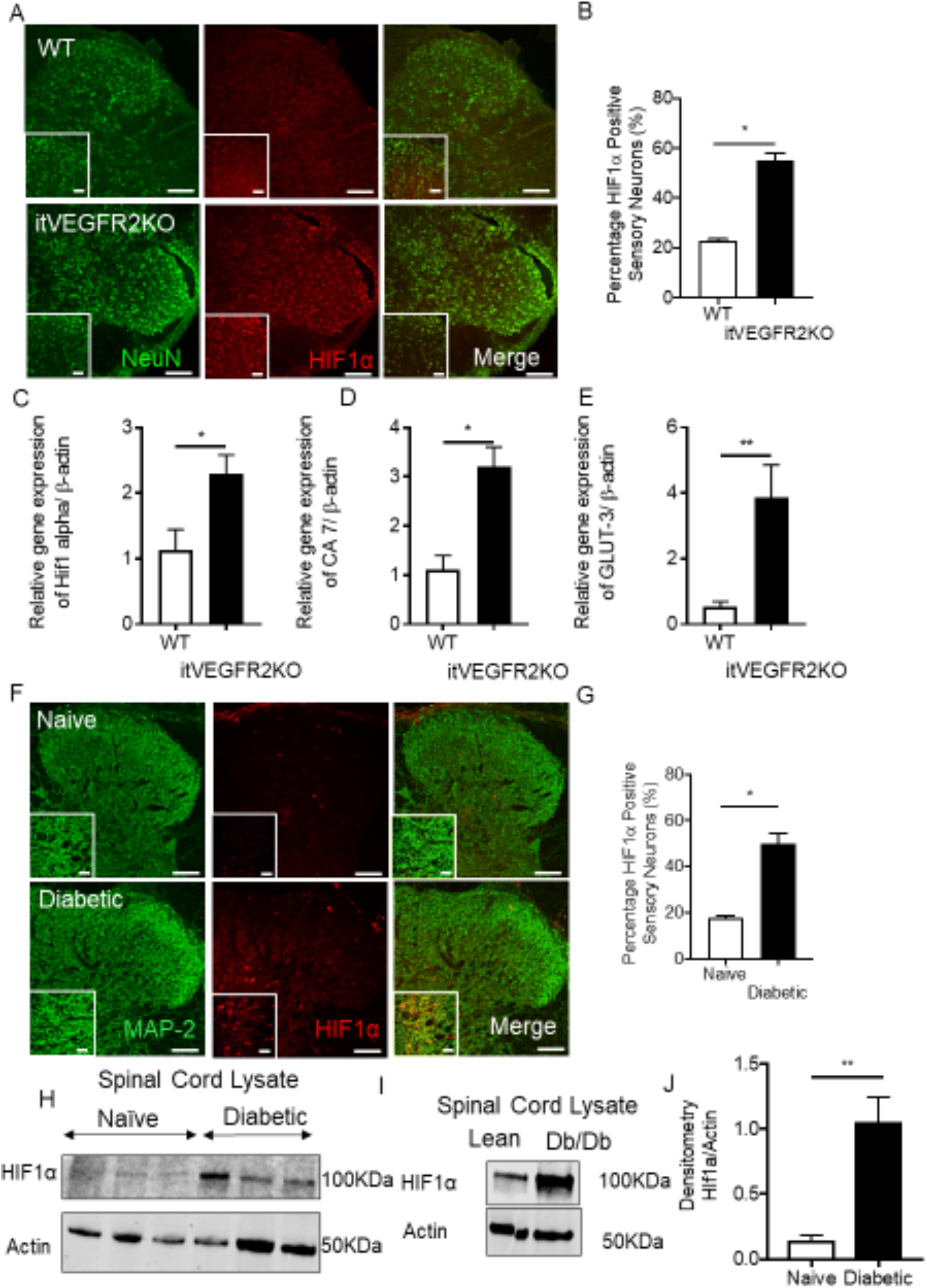
Spinal cord vasculopathy induces HIF1*α* expression in neurons in the dorsal horn [A] 8 days after OHT treatment HIF1*α* protein immunoreactivity was increased in sensory neurons of the dorsal horn in VEGFR2^scECKO^ mice (Representative confocal images of WT and VEGFR2^scECKO^ co-labelled with NeuN, low power images (main) scale bar = 100μm, high power images (inset) = 50 μm). [B] There was an increase in the percentage of dorsal horn sensory neurons expressing HIF1*α* in VEGFR2^scECKO^ (*P<0.05, Unpaired T Test, n= 5 per group). mRNA quantification from lumbar spinal cord samples demonstrated increased expression of [C] HIF1*α* (*P<0.05, Unpaired T Test, n= 5 per group), [D] CA7 (*P<0.05, Unpaired T Test, n= 5 per group) and [E] GLUT3 (**P<0.01, Unpaired T Test, n= 5 per group) in VEGFR2^scECKO^ mice. [F] In a rat model of type 1 diabetes and age matched controls HIF1*α* protein immunoreactivity was determined in the dorsal horn neurons (Representative confocal images of naïve and diabetic co-labelled with MAP2, low power images scale bar = 200μm, high power images = 100μm). [G] There was an increased percentage of dorsal horn neurons expressing HIF1*α* in diabetic rats when compared with age-matched naïve control animals (*P<0.05, Unpaired T Test, n= 5 per group). [H] representative western blots showing an increase HIF1*α* protein expression in lumbar spinal cord protein lysates extracted from types 1 (STZ induced) diabetic rats [I] Densitometry quantification demonstrated an increase HIF1*α* protein expression (**P<0.05, Unpaired T Test, n= 5 per group) [I] Immunoblot of protein from spinal cord of type 2 diabetic mouse (db/db) compared with controls.

**Figure 6 –.**
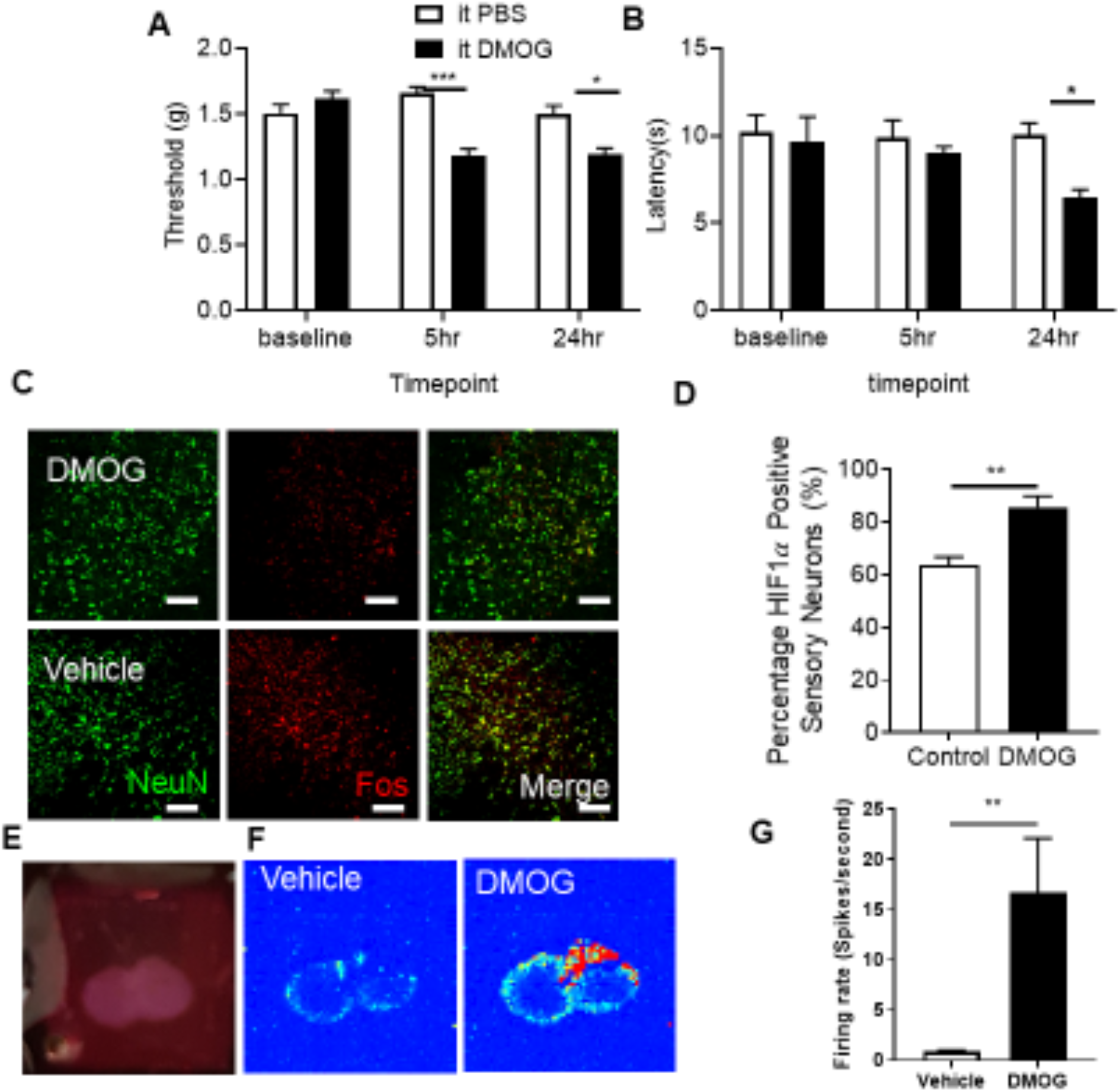
Chemically induced hypoxia causes sensory neuronal activation in dorsal horn and initiation of pain hypersensitivity Intrathecal injection of 1mM DMOG in C57Bl6 male mice led to [A] lower mechanical withdrawal thresholds at 5 and 24 hrs as well as [B] reductions in heat withdrawal latency when compared with vehicle treated mice (*P<0.05, ***P<0.001 Two Way ANOVA with Tukey’s multiple comparison, n=10 per group). Fos, a marker of neuron activation, was increased in dorsal horn neurons ([C] (co labelled with neuronal marker NeuN, scale bar = 50 μm) following intrathecal DMOG injection (**P<0.01, Unpaired T Test, n=5 per group). [E] Neuronal activity was recorded in vitro from lumbar spinal cord slices using multi-electrode arrays and [F] basal neuronal activity heat maps 24 hours following vehicle and DMOG treated samples generated. [G] DMOG treated lumbar spinal cord slices demonstrate increased basal neuronal activity compared to vehicle treated samples (**P<0.01, Unpaired T Test, n=3).

### Hypoxia induction <u>and</u> Onset <u>of</u> Chronic Pain

These results all suggest that hypoxia due to vascular loss can induce behavioural hypersensitivity. To test this, the chemical hypoxia mimetic dimethyloxalylglycine (DMOG) was intrathecally injected into WT mice. 24 hours after intrathecal injection of DMOG there was a significant reduction in mechanical withdrawal threshold (Fig. 6A) and a decrease in heat withdrawal latency (Fig. 6B) compared with vehicle treatment. CA7 was found to be expressed in the spinal cord neurons alongside GLUT3 and HIF1*α*, with expression associated with excitatory sensory neurons expressing TACR1 (Supplemental Figure 6C). Additionally, there was an increased number of dorsal horn sensory neurons expressing fos immunoreactivity following intrathecal DMOG injection (Fig. 6C and 6D). Furthermore, isolated spinal cord neurons presented increased potassium chloride induced calcium influx following 24hr treatment to hypoxia mimetic (DMOG) when compared with vehicle treated samples (Supplemental Figure 6D). Using multi-electrode arrays on spinal cord splices (Fig. 6E) we showed increased baseline neuronal activity after DMOG treatment in the dorsal horn aspect (Fig. 6F) compared with vehicle treated lumbar spinal cord slices (Fig. 6G).

### Carbonic Anhydrase Mediated Hypoxia Induced Chronic Pain

We identified that CA7 was upregulated following induction of hypoxia by VEGFR2 knockout in spinal cord endothelial cells. To determine whether this mediated neuronal hypersensitivity we reproduced this by treatment with DMOG for 24hrs, which showed increased protein expression (Fig. 7A representative western blot) of HIF1*α* (Fig. 7B) and CA7 (Fig. 7B) compared with vehicle treatment. Similarly, intrathecally administered DMOG led to increased protein expression (Fig. 7D representative western blot) of HIF1*α* (Fig. 7E) and CA7 (Fig. 7F). Acetazolamide (ACZ) a non-selective carbonic anhydrase inhibitor delivered via intraperitoneal injection attenuated DMOG induced nociceptive behavioural hypersensitivity. ACZ increased mechanical withdrawal thresholds returning to comparable thresholds to the control group (Fig. 7G). Similarly, heat withdrawal latencies were increased to normal values in the DMOG treated experimental cohort (Fig. 7H). ACZ also increased withdrawal latencies in VEGFR2^scECKO^ mice, partially restoring them towards vehicle treatment (Fig. 7I).

**Figure 7 –.**
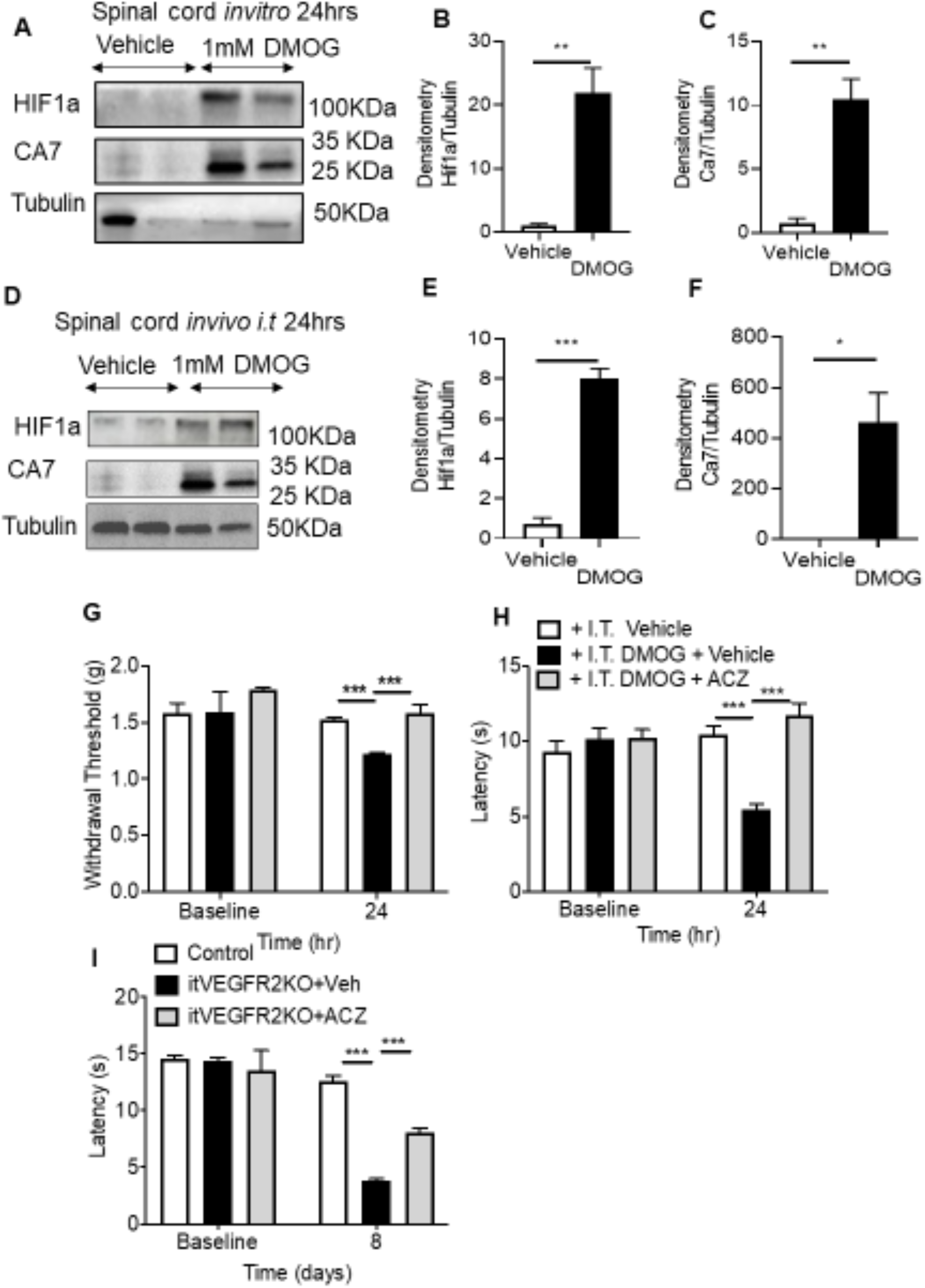
Inhibition of Carbonic Anhydrase 7 attenuates hypoxia induced pain hypersensitivity [A] *In vitro* lumbar spinal cord slices were treated with either Vehicle or 1mM DMOG for 24hrs to determine protein expression of HIF1*α* and CA7 (Representative western blots), with [B] DMOG treatment increasing expression of [B] HIF1*α* and [C] CA7 compared with vehicle treatment (**P<0.01, Unpaired T Test, n=5 per group). In C57Bl6 mice, lumbar spinal cord protein lysate samples ([D] Representative western blots) 24hrs following intrathecal delivery of 1mM DMOG demonstrated an increase in protein expression of [E] HIF1*α* and [F] CA7 compared with vehicle treated animals (*P<0.05, ***P<0.001, Unpaired T Test, n=3 per group). [G] Intraperitoneal treatment of acetazolamide (ACZ) led to attenuation of DMOG induced reduction in mechanical withdrawal thresholds and [H] heat withdrawal latencies (**P<0.01, ***P<0.001 Two Way ANOVA with Tukey’s multiple comparison, n=5 per group). ACZ treatment led to nociceptive withdrawal thresholds returning to baseline values and comparable to vehicle treated animals. [I] Intraperitoneal ACZ inhibited heat hypersensitivity in VEGFR2^scECKO^ mice with heat withdrawal latencies increased compared with VEGFR2^scECKO^ mice treated with vehicle (***P<0.001 Two Way ANOVA with Tukey’s multiple comparison, n=5 per group).

## Discussion

Chronic pain is induced through alterations in sensory neuronal activity, with the vascular system playing a key role in the manifestation of long-term pain hypersensitivity through induction of pro-inflammatory signalling(31). Here we have explored how vascular dysfunction through curtailed blood flow and subsequent hypoxia at the level of the spinal cord could modulate nociceptive processing. Currently those mechanisms associated with induction of central sensitisation are fundamental to the development of neuropathic pain, with chloride and bicarbonate homeostasis integral to GABA mediated disinhibition. In this study we identify that reduced blood perfusion of the spinal cord induces hypoxia mediated activation of dorsal horn spinal sensory neurons and chronic pain manifestation. Importantly chronic pain behaviours were suppressed by inhibition of carbonic anhydrase activity.

### Vasculopathy, VEGFR2 signalling and Nociception

Impairment in nervous system blood flow is highly associated with the development of neurological disease e.g. Alzheimer’s, dementia (14, 32), whilst also a fundamental cause of ischaemic disease of the nervous system such as stroke(33). The underlying mechanisms by which the somatosensory system residing in the central nervous system adapts to such fluctuations in the microenvironment remain unclear. Investigations presented here suggest that a reduction of the vascular supply to the spinal cord is sufficient to initiate pain.

Chronic pain is a highly prevalent pathological condition, affecting large proportions of the population, with the onset and severity correlated with progression in age and/or disease (1). Incidentally capillary health is also greatly influenced by age and disease, with a decline in vessel number and increased frequency of vasoconstrictive tone associated with neurological pathology(14, 34). Endothelial degeneration in the spinal cord capillary network is implicated in the development of neuropathic pain(17), with loss of endothelial specific VEGFR2 signalling highly attributable to the manifestation of pain hypersensitivity. Vascular endothelial growth factor 2 (VEGFR2) signalling is integral for the development and maintenance of vascular function in health and disease through maturation of the capillary network (35) and regulation of vessel permeability (36, 37). In neurological disease, alterations in VEGFR2 signalling results in vascular defects and consequently a decline neuronal function(38-40). Furthermore, supplementation with VEGF-A has been shown to prevent the development of nervous system vasculopathy leading to the suppression of neurological disease (17, 18, 40). In relation to the spinal cord, VEGFR2 expression and activity was attributable to spinal cord endothelium function via manipulation of vascular permeability. Consequently, we showed that the loss of VEGFR2 expression in the spinal cord endothelium led to a reduction in the capillary network throughout the dorsal horn, supporting the integral role of VEGFR2 in maintaining capillary health. Previous work (17) had explored systemic VEGFR2 knockdown, plausibly influencing a wider context in relation to the somatosensory network (41). Here in this study VEGFR2 knockdown was restricted to the spinal cord endothelium, enabling interrogation of the interplay between the vascular and neuronal systems in relation to pain modulation. Induction of VEGFR2 knockdown in specifically the spinal cord endothelium leads to thermal nociceptive hypersensitivity.

### Hypoxia induced Chronic Pain

The loss of spinal cord microvasculature in the dorsal horn of the spinal cord and subsequent loss of capillary perfusion was associated with the induction of hypoxia signalling in rodent models of chronic pain. Neuronal tissue is highly metabolically active therefore an established blood supply to neural tissues is pivotal, with neuronal activity coupled to provision of nutritional support(42). As a consequence of this, neurons are highly sensitive to fluctuations in oxygen and nutrient delivery (43, 44), with neuronal integrity compromised by diminished nutritional support. Changes in the neural tissue environment such as diminished tissue oxygen level are detrimental to neural tissue integrity, altering neuronal activity leading to pathological neurological disabilities, seizures and cognitive impairments (45). Following induction of vascular degeneration and induction of hypoxia mechanisms, sensory neurons in the dorsal horn increased levels of neuronal activity. This activity has been observed in rodent models of chronic pain (46), and in spinal cord damage observed in diabetic neuropathic pain patients (47, 48).

Impaired blood flow to the peripheral sensory nervous system induces exacerbated pain behaviours in rodents that is accompanied by enhanced peripheral sensory activity (49, 50) typifying onset of chronic pain states following vascular occlusion. Currently there are few published investigations into the relationship between the sensory neuron at the level of the spinal cord and reduced blood flow. However, reductions in blood flow and consequently tissue oxygen level impairs motor neuron activity, underlying the significance of blood flow to neuron function (34). In this study using rodent models of vasculopathy that induce chronic pain we demonstrated pronounced hypoxia signalling in the dorsal horn of the spinal cord, in association with neuronal activation. Increased expression of C-fos (marker of neuronal activation) was accompanied by elevated expression levels of hypoxia markers hypoxia inducible factor 1 *α* (HIF1*α*), glucose transporter 3 (GLUT3) and carbonic anhydrase 7 (CA7). Previously exploration of the impact of hypoxia upon the sensory neuron demonstrated the dorsal root ganglia sensory neuron is activated under such low oxygen conditions (51, 52). Furthermore, development of diabetic neuropathic pain is inherently associated with a neurovascular disruption with reductions in oxygen attributable to sensory neurodegeneration(53). Sensory neuronal activity has previously been demonstrated to be influenced by hypoxia and HIF1*α* activity (51). Reductions of HIF1*α* expression in dorsal root ganglia sensory neurons was associated with a decline in the number of c-fos expressing neurons following peripheral sensory neuron activation (51). However, hypoxia induced expression of neuron specific GLUT3 and CAR7 highlights a putative adaptation of the dorsal horn sensory neuron to alterations in the tissue microenvironment, underlying plausible avenues by which synaptic plasticity may evolve to underlie chronic pain manifestation. Of interest, we saw no change in DRG vessel markers in the VEGFR2^scECKO^, and yet changes in pain behaviour were seen, suggesting that while DRG may contribute, there is also significant spinal cord involvement.

Identifying those neuronal adaptation mechanisms responsible for modulation of neuronal function and synaptic plasticity are integral in the understanding how chronic pain develops (10, 54). Dysregulation of GABAergic signalling is widely recognised in the development of chronic pain states through differential regulation of chloride, potassium and bicarbonate homeostasis (55). Crucially loss of potassium chloride cotransporter 2 (KCC2) in the spinal cord leads to the onset of chronic pain and sensory neuronal excitation. KCC2 inhibition or downregulation (such as in times of pathological pain and hypoxia reduced KCC2 activity) leads to accumulation of intracellular chloride. Subsequent activation of GABA receptor A (GABA^A^) leads to efflux of chloride leading to excitation of the sensory neuron. However, recent evidence identifies that regulation in the transport of chloride and bicarbonate influences spinal cord neuronal activation and chronic pain states(56). GABA^A^ not only induces chloride efflux but bicarbonate is also transported out of the neuron. Consequently, suppression of bicarbonate synthesis via carbonic anhydrase activity through administration of acetazolamide (ACZ) inhibits sensory neuron activity and nociception at the level of the spinal cord (11, 57, 58). Similarly, our work shows that hypoxia induced pain can be ameliorated by ACZ treatment, which in part is driven by hypoxia induced upregulation of CA7 expression. CA7 is a cytoplasmic isoform that utilises carbon dioxide and water to synthesise bicarbonate. In relation to neuronal function it has been established that in the hippocampus that GABA^A^ mediated neural excitation is dependent upon intracellular bicarbonate levels maintained via cytoplasmic carbonic anhydrase activity (59), with CA7 associated with aberrant hippocampal neural activity responsible for seizures (60). Furthermore, CA7 is expressed in the spinal cord (60) in predominantly excitatory sensory neurons, along with membrane bound CA12 (61, 62). However, a CA7 knockout mouse demonstrated no alterations in nociceptive behaviour in comparison with a control mouse (56). Of note, that study explored naïve animals and our results highlight that CA7 is expressed at low levels in such an instance. It is plausible that induction of GABA disinhibition such as in a pathological pain state driven by hypoxia is needed to induce CA7 dependent sensory neural excitation.

This study presents a novel mechanism that activates excitatory sensory neuron networks in the spinal cord to initiate chronic pain. By understanding the intrinsic plasticity of the sensory neural network in relation to the impact of specific disease states upon the somatosensory nervous system enables greatly informed development of novel analgesics therapies.

## Acknowledgments

RPH, MD, NW, RM, LW and SB performed the experimental work. RPH, MD, NW, RM, LW and SB, CS LFD & DOB contributed to the conception or design of the work in addition to acquisition, analysis or interpretation of data for the work. RPH, MD, NW, RM, LW and SB, LFD & DOB drafted the work or revised it critically for important intellectual content. RPH drafted the manuscript with contributions from all authors. All authors approved the final version of the manuscript, agree to be accountable for all aspects of the work in ensuring that questions related to the accuracy or integrity of any part of the work are appropriately investigated and resolved all persons designated as authors qualify for authorship, and all those who qualify for authorship are listed.

## Supplemental Figures

**Supplemental Figure 1 –.**
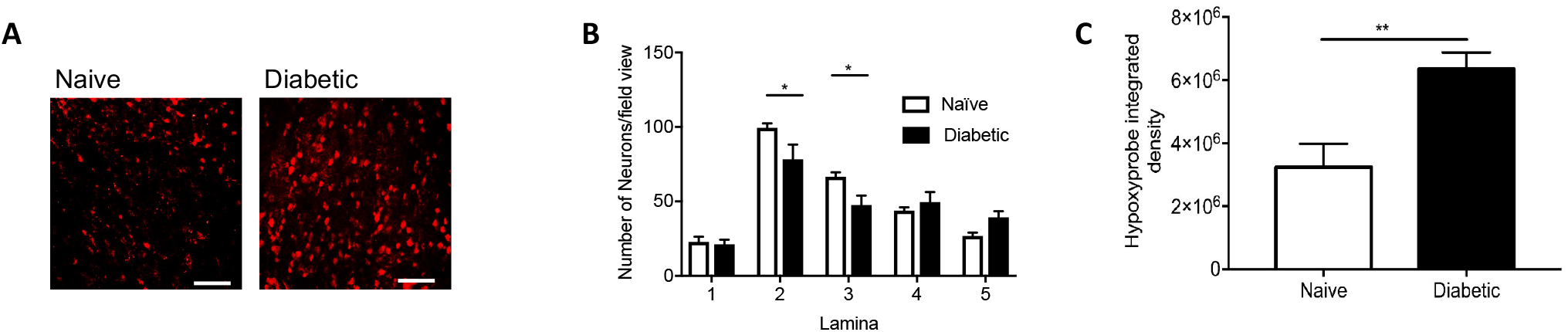
<u>Diabetes induced hypoxia in the dorsal horn</u> [A] Confocal microscopy of the dorsal horn was used to identify hypoxyprobe (HP) labelled sensory neurons (high power images scale bar = 50 μm) in Naïve and Diabetic Sprague Dawley female rats. [B] There were decreased neuronal number in the dorsal horn of the spinal cord in diabetic Sprague Dawley female rats. [C] In Diabetic Sprague Dawley female rats demonstrated increased level hypoxyprobe fluorescence compared to naive control animals (**P<0.01, Unpaired T Test, n=5 per group).

**Supplemental Figure 2 –.**
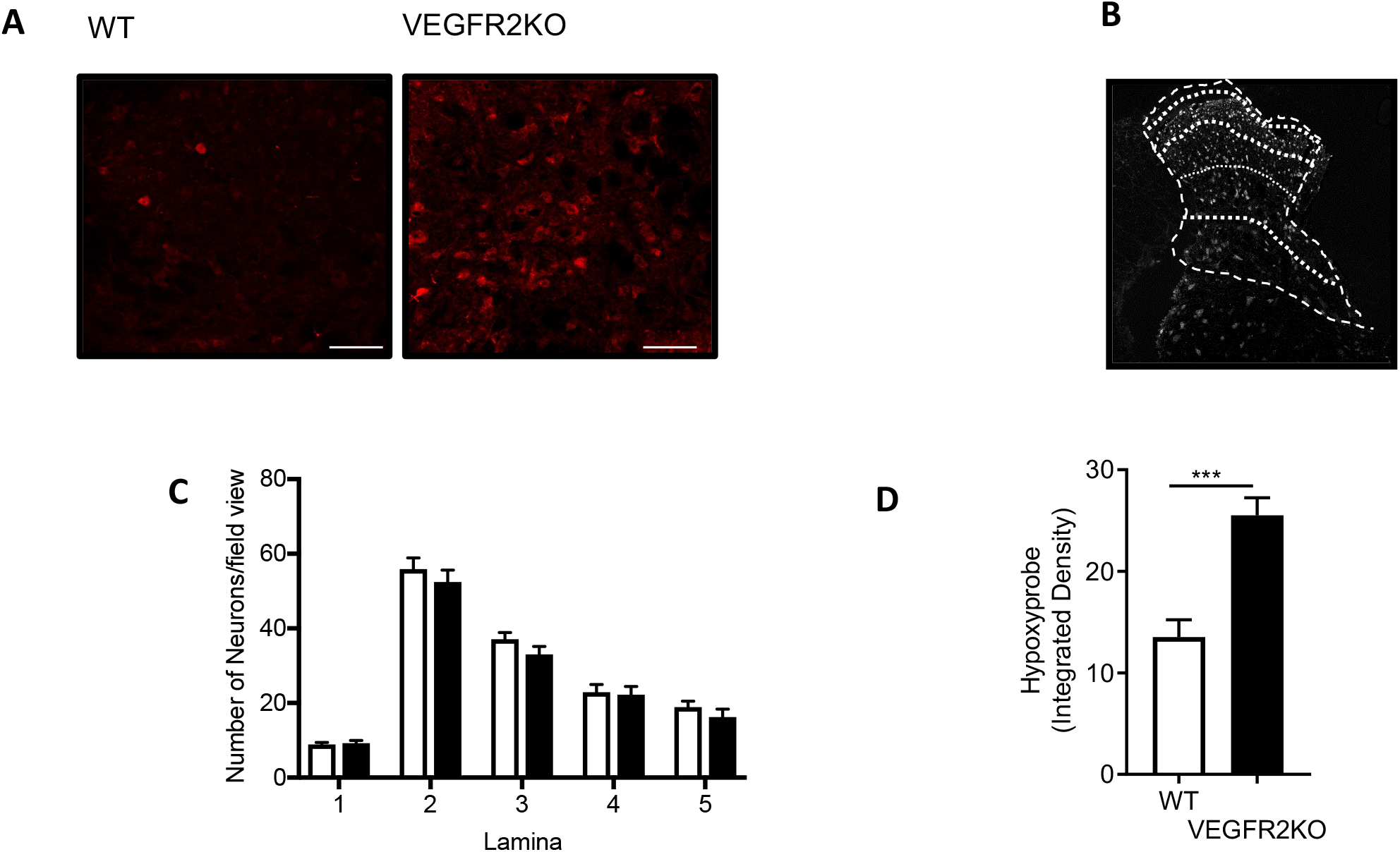
Endothelial *VEGFR2Knock-Outinducedhypoxiainthedorsalhorn* [A] Confocal microscopy of the dorsal horn was used to identify hypoxyprobe (HP) labelled sensory neurons (high power images scale bar = 50 μm) in WT and VEGFR2^ECKO^ mice. [B] Diagrammatic breakdown of the lamina of the dorsal horn to provide quantification of the neuron number per lamina. [C] There were no alterations in neuronal number in the dorsal horn of the spinal cord in either the WT or VEGFR2^ECKO^ mice (white bars=WT, black bars = VEGFR2^ECKO^). [D] In VEGFR2^ECKO^ mice demonstrated increased level of hypoxyprobe fluorescence compared to WT control animals (*P<0.05, Unpaired T Test, n=5 per group).

**Supplemental Figure 3 –.**
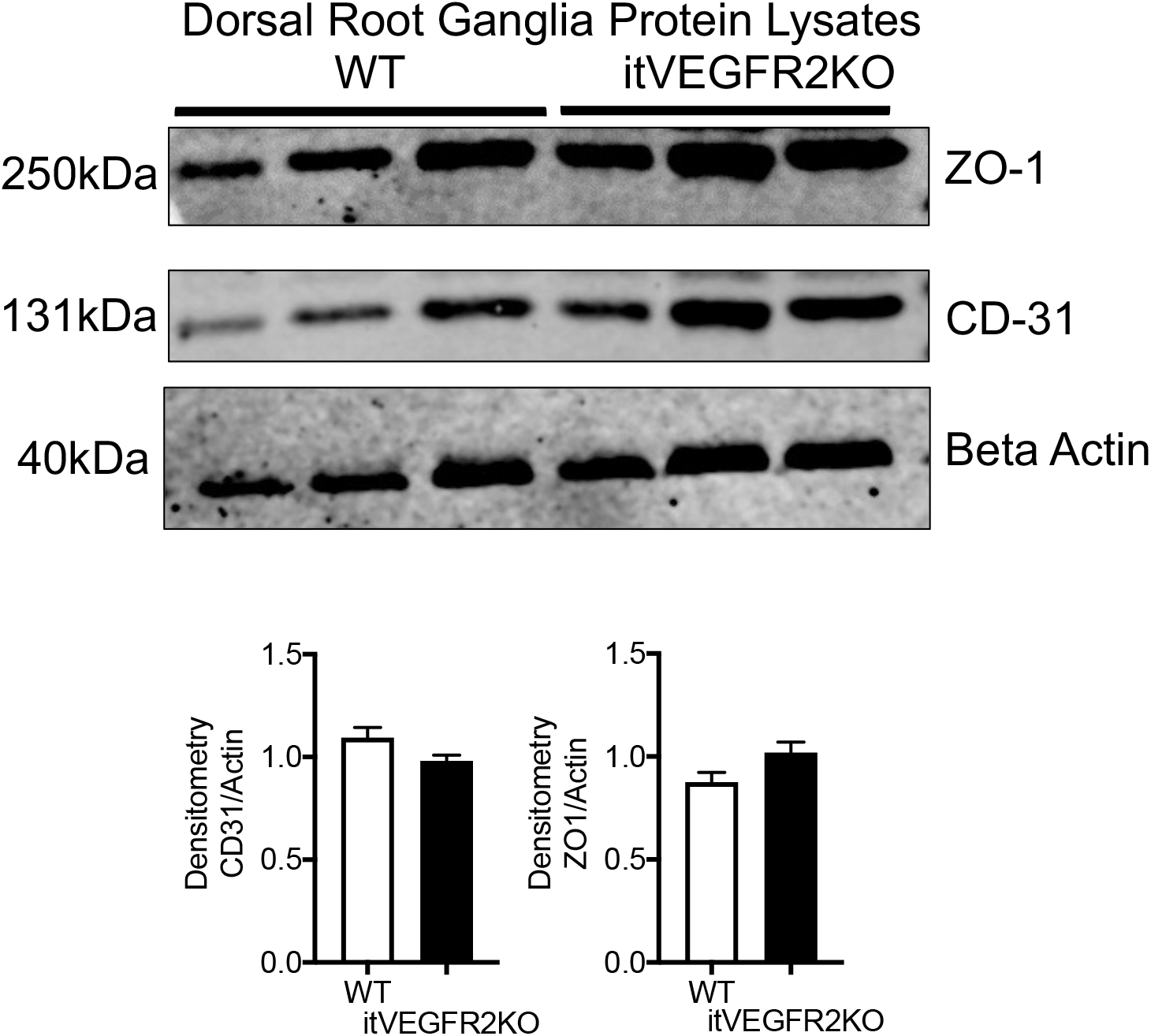
<u>DRG vasculature was unaffected by intrathecal OHT injection</u> OHT treatment led to no alterations in protein expression of endothelial markers in the dorsal root ganglion extracted from WT or itVEGFR2 mice. Densitometry quantification of protein samples extracted from dorsal root ganglion (pooled dorsal root ganglion from lumbar 3, 4 and 5) demonstrated no change in ZO1 or CD31 expression in either WT or itVEGFR2KO mice.

**Supplemental Figure 4 –.**
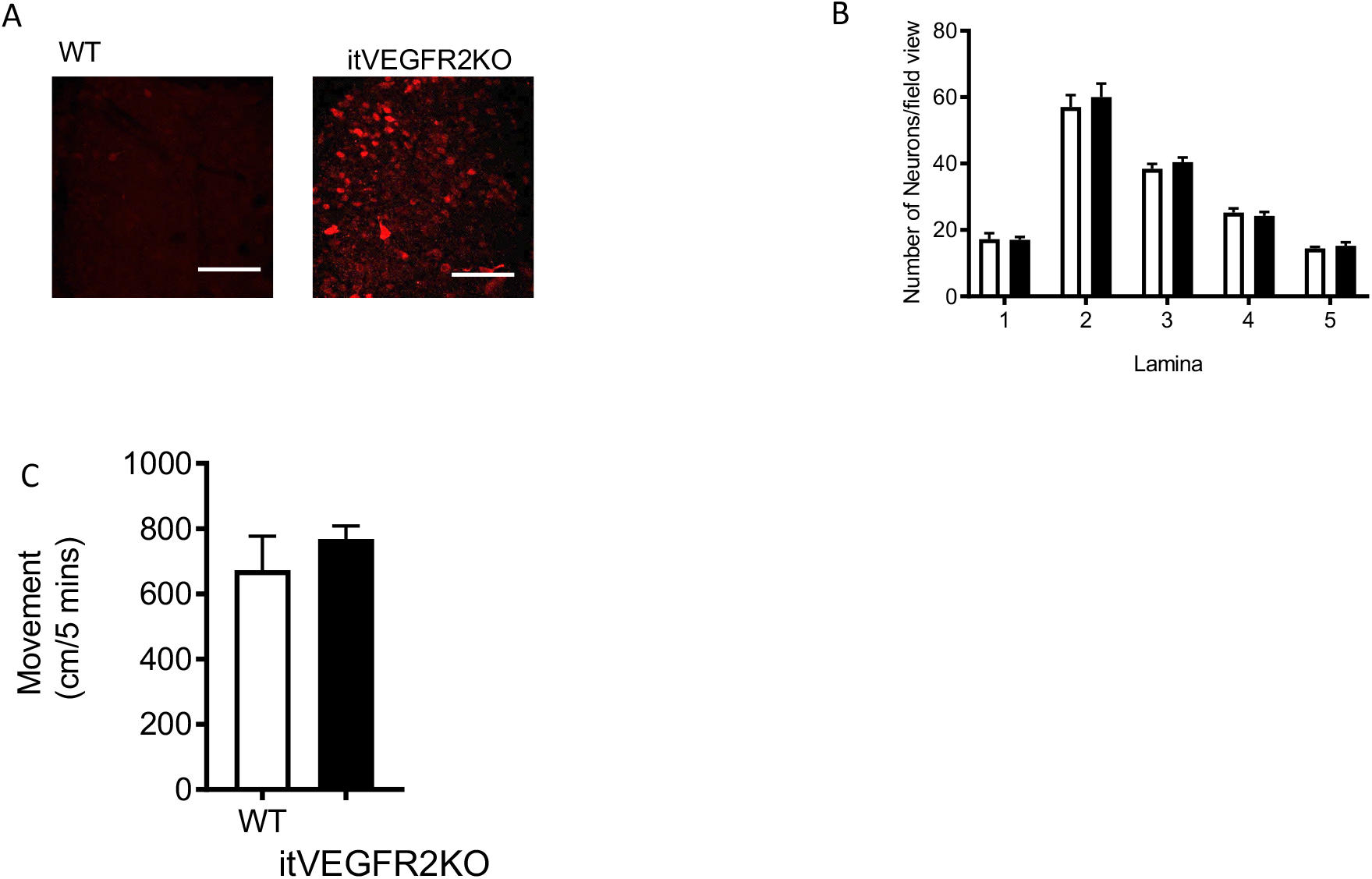
<u>Spinal cord vasculopathy induces hypoxia and no impact upon motor behaviour</u> [A] Representative confocal images of Hypoxyprobe labelled sensory neurons in the dorsal horn of WT and VEGFR2^scECKO^ mice (high power images scale bar = 50 μm). [B] There were no alterations in neuronal number in the dorsal horn of the spinal cord in either the WT or VEGFR2^scECKO^ mice (white bars=WT, black bars = VEGFR2KO). [C] There were no alterations in motor behavior with WT and VEGFR2^scECKO^ mice moving the same distance in the testing environment.

**Supplemental Figure 5 –.**
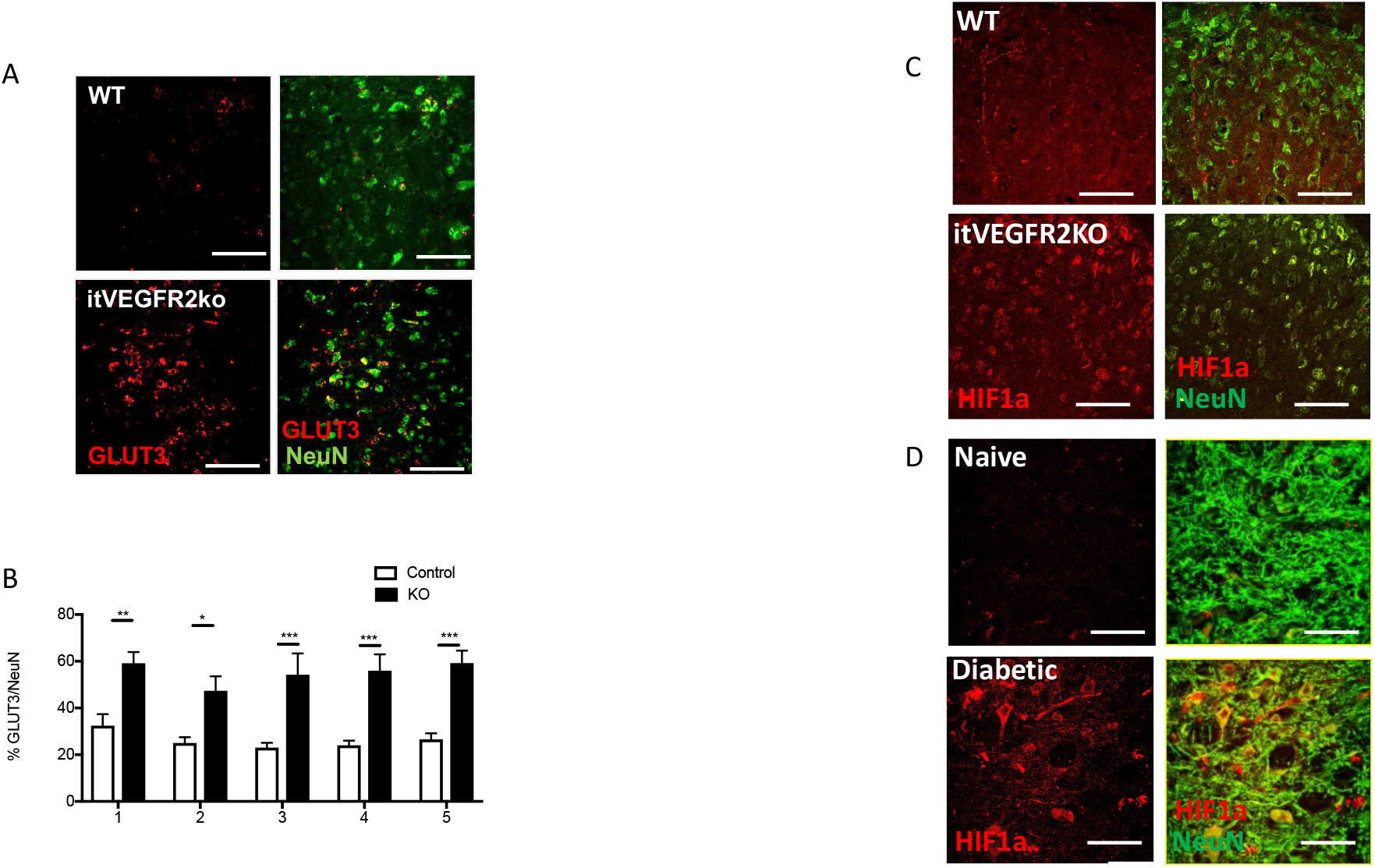
<u>Spinal cord vasculopathy induces GLUT3 and HIF1*α* expression in sensory neurons in the dorsal horn</u> [A] 8 days following OHT treatment GLUT3 protein immunoreactivity was increased in sensory neurons of the dorsal horn in VEGFR2^scECKO^ mice (Representative confocal images of WT and VEGFR2^scECKO^ co-labelled with NeuN, high power images scale bar = 50 μm). [B] There was an increase in the percentage of dorsal horn sensory neurons expressing GLUT3 in VEGFR2^scECKO^ mice when compared to WT mice (*P<0.05, **P<0.01, ***P<0.001 Two way ANOVA with post Bonferroni test, n= 5 per group). [C] 8 days following OHT treatment HIF1*α* protein immunoreactivity was increased in sensory neurons of the dorsal horn in VEGFR2^scECKO^ mice (Representative confocal images of WT and VEGFR2^scECKO^ co-labelled with NeuN, high power images scale bar = 50 μm). [D] HIF1*α* protein immunoreactivity was increased in sensory neurons of the dorsal horn in STZ induced diabetic Sprague Dawley rats versus age matched control rats (Representative confocal images of WT and VEGFR2^scECKO^ co-labelled with NeuN, high power images scale bar = 50 μm).

**Supplemental Figure 6 –.**
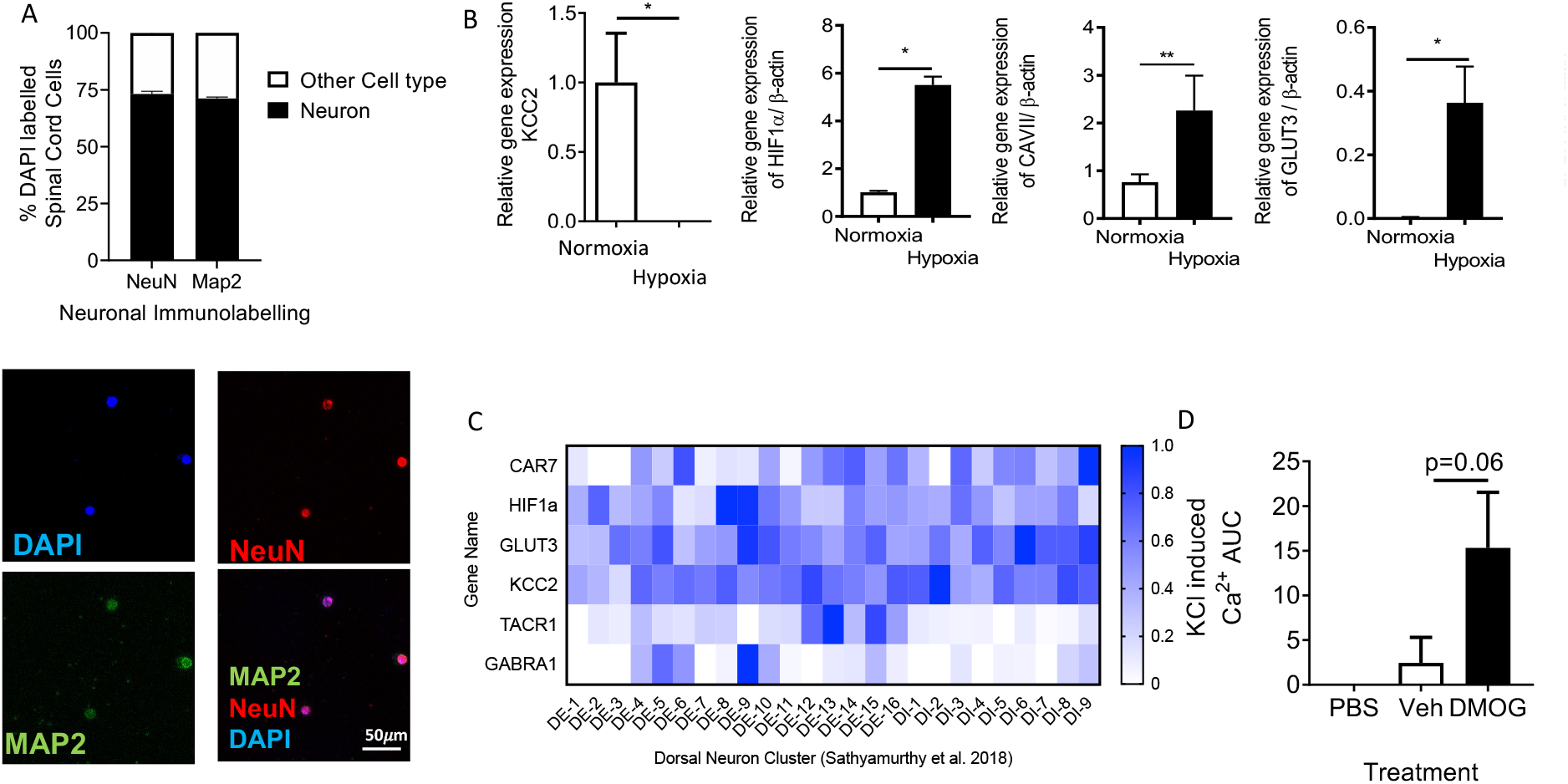
<u>Hypoxia induced CA7, GLUT3 and HIF1*α* expression in the spinal cord</u> [A] Lumbar spinal cord neurons were isolated from C57bl6 mice (labelled with neuronal markers NeuN and MAP2 as well as nuclei marker DAPI; scale bar = 50μm). [B] Isolated neurons were exposed to normoxic and hypoxic conditions. Hypoxia led to the increased expression of induced CA7, GLUT3 and HIF1*α*, as well as reduced expression of KCC2 (*P<0.05, **P<0.01, Unpaired T Test). [C] Additionally, CA7 was identified in spinal cord neurons with particular association with excitatory sensory neurons (TACR1 positive) (GEO dataset = Sathyamurthy et al. 2018). [D] Isolated spinal cord neurons treated (24hrs) with either chemical induction of hypoxia through 1mM DMOG treatment versus vehicle control demonstrated increased neuronal response following stimulation with KCL (p=0.06 One Way ANOVA).

